# Rapid GPR183-mediated recruitment of eosinophils to the lung after *Mycobacterium tuberculosis* infection

**DOI:** 10.1101/2022.02.18.480919

**Authors:** Andrea C. Bohrer, Ehydel Castro, Claire E. Tocheny, Maike Assmann, Benjamin Schwarz, Eric Bohrnsen, Michelle A. Makiya, Fanny Legrand, Kerry L. Hilligan, Paul J. Baker, Flor Torres-Juarez, Zhidong Hu, Hui Ma, Lin Wang, Liangfei Niu, Wen Zilu, Sang H. Lee, Olena Kamenyeva, Tuberculosis Imaging Program, Keith D. Kauffman, Michele Donato, Alan Sher, Daniel L. Barber, Laura E. Via, Thomas J. Scriba, Purvesh Khatri, Yanzheng Song, Ka-Wing Wong, Catharine M. Bosio, Amy D. Klion, Katrin D. Mayer-Barber

**Author notes:** These authors contributed equally and are listed in alphabetical order (Andrea C. Bohrer, Ehydel Castro).

## Abstract

Influx of eosinophils into the lungs is typically associated with type-II responses during allergy, fungal and parasitic infections. However, we previously reported that eosinophils accumulate in lung lesions during type-I inflammatory responses to *Mycobacterium tuberculosis* (Mtb) in humans, macaques, and mice where they contribute to host resistance. Here we show eosinophils migrate into the lungs of macaques and mice as early as one week after Mtb-exposure. In mice this influx was CCR3 independent and instead required cell-intrinsic expression of the oxysterol-receptor GPR183, which is highly expressed on human and macaque eosinophils. Murine eosinophils interacted directly with bacilli-laden alveolar macrophages, which upregulated the oxysterol-synthesizing enzyme Ch25h, and eosinophil recruitment was impaired in Ch25h deficient mice. Our findings show that eosinophils are among the first cells from circulation to sense and respond to Mtb infection of alveolar macrophages and reveal a novel role for GPR183 in the migration of eosinophils into lung tissue.

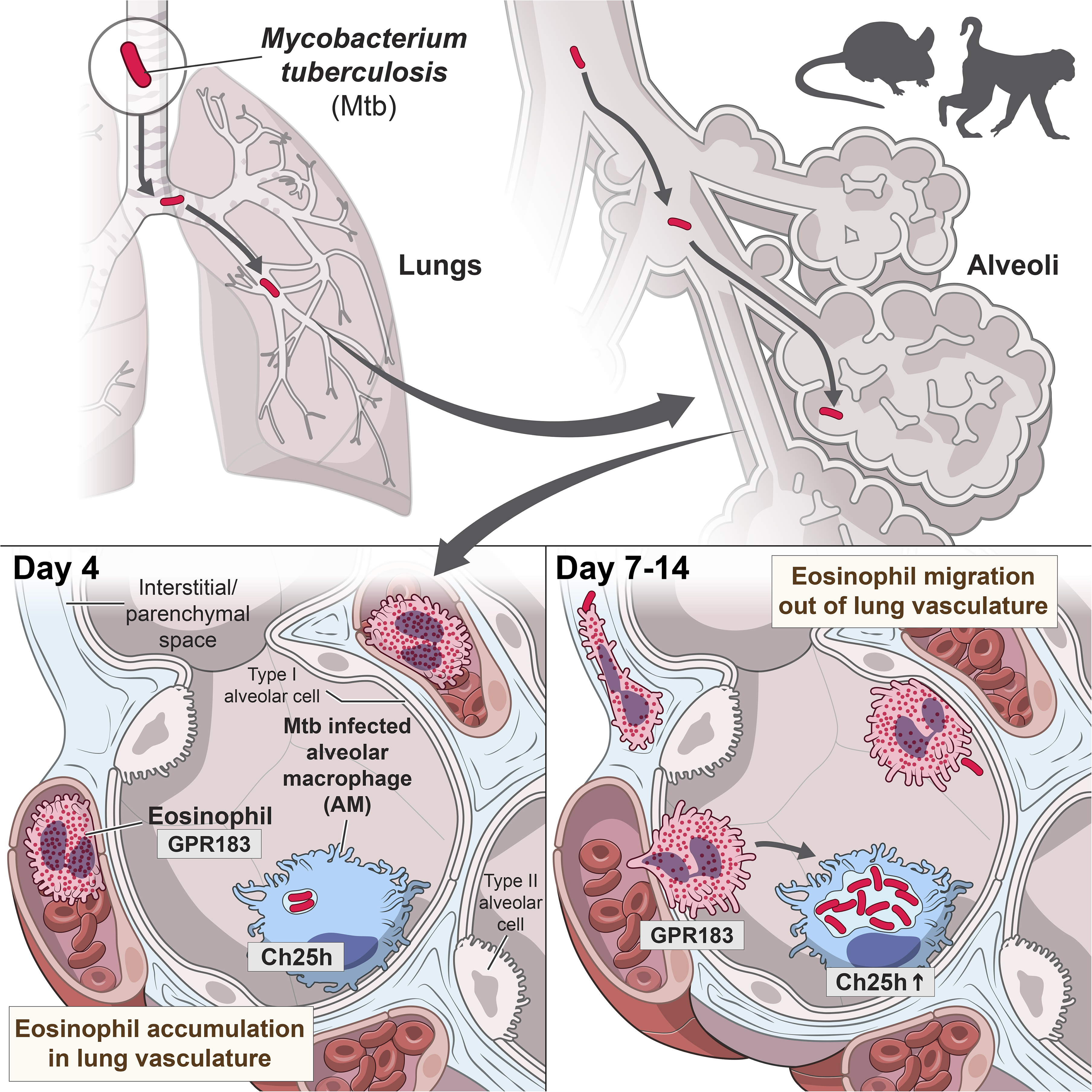

**HIGHLIGHTS:**

In mice and macaques eosinophils accumulate early in Mtb-infected lungs preceding neutrophils Eosinophils interact with Mtb-infected cells in the alveoli in mice
Early pulmonary eosinophil migration occurs independently of CCR3 in mice
Early lung migration in mice requires Ch25h and eosinophil-intrinsic GPR183 expression

## INTRODUCTION

*Mycobacterium tuberculosis* (Mtb) is the intracellular bacterial pathogen responsible for tuberculosis (TB) disease, which is a leading cause of mortality world-wide. Host resistance during Mtb infection depends on T helper type-I responses associated with IFN-γ and IL-12 pathways, hallmark features of canonical type-I immunity (Ernst, 2012; Kinsella et al., 2021). Infection with Mtb is established when inhaled bacteria infect lung airway resident alveolar macrophages (AM) (Cohen et al., 2018; Ernst, 2012; Rothchild et al., 2019). The subsequent innate cellular response that immediately follows Mtb infection of AM and precedes T helper cell involvement is not well understood *in vivo* and is an area of active investigation (Corleis and Dorhoi, 2020; O’Garra et al., 2013; Philips and Ernst, 2012; Ravesloot-Chavez et al., 2021). Early host responses in the lungs can influence long-term disease trajectory by affecting cell recruitment, adaptive immunity, granuloma formation, bacterial dissemination, disease tolerance and progression (Cadena et al., 2016). However, very little is known about the specific innate immune cell types responding to or interacting with infected AM during the first two weeks after infection. Granulocytes, in particular neutrophils, are often the first responders to microbial invasion. For example, during acute bacterial pneumonia the rapid influx of granulocytic phagocytes is critical for infection control (Prince, 2013; Yang et al., 2012). Accordingly, neutrophils are generally thought to be among the first cells to respond to Mtb-infected AM (Blomgran et al., 2012; Corleis and Dorhoi, 2020). However, at later stages of chronic Mtb infection, neutrophils have been shown to provide a replicative niche for Mtb bacilli and recruitment of neutrophils to the lung correlates with TB disease progression and increased bacterial loads (Eum et al., 2010; Lowe et al., 2013; Lyadova, 2017; Mishra et al., 2017; Muefong and Sutherland, 2020; Scott et al., 2020).

In contrast to neutrophils, eosinophils are key innate effector cells in type-II inflammatory settings (Klion and Nutman, 2004; Simon et al., 2020). Consequently, *in vivo* lung eosinophil responses have primarily been studied in experimental models of allergy, parasitic helminth infections (Klion et al., 2020; Travers and Rothenberg, 2015), viral infections that exhibit some aspects of type-II immunity (i.e. respiratory syncytial virus) or fungal infections with *Aspergillus* and *Cryptococcus* (Lilly et al., 2014; Phipps et al., 2007). More recently it has become clear that eosinophils take on diverse and complex roles in lung tissue homeostasis, barrier function and wound healing responses (Rosenberg et al., 2013; Shah et al., 2020). In this context, we have recently reported that eosinophils are enriched in lung lesions, but not in circulation, in patients with active TB disease and that they are present and functionally activated in granulomas of *Mtb*-infected rhesus macaques (Bohrer et al., 2021). Moreover, we showed that mouse strains genetically deficient in eosinophils displayed greater susceptibility to tubercular disease including decreased survival and increased bacterial loads (Bohrer et al., 2021). These previous findings suggest that eosinophils participate in the innate immune response to tuberculosis. However, the mechanism of eosinophil recruitment into the lungs during the highly type-1 polarized inflammation of Mtb infection remains unclear.

Here we show that eosinophils are very rapidly recruited from circulation into the lungs of both mice and rhesus macaques after exposure to Mtb, and eosinophils can be visualized interacting with Mtb-infected cells in mice. Pulmonary eosinophil recruitment in mice occurs independently of CCR3, the canonical chemokine receptor for eosinophil migration during type-II inflammation. Instead, we show that in mice optimal recruitment of circulating eosinophils to Mtb-infected lungs requires cell-intrinsic oxysterol sensing through GPR183. Moreover, infected murine alveolar macrophages selectively upregulate the oxysterol-producing enzyme Ch25h, and pulmonary recruitment of eosinophils is reduced in the absence of Ch25h. Thus, our findings reveal that GPR183-dependent oxysterol sensing is a novel mediator of eosinophil recruitment to the lungs and acts as an early alarm signaling the presence of Mtb infection in the airways to cells in circulation.

## RESULTS

### Eosinophils dominate the early granulocyte response in the airways of Mtb-infected rhesus macaques

Our previous work demonstrated the presence of eosinophils in established TB granulomas of nonhuman primates (NHP) two-three months after Mtb infection and the importance of eosinophils for host-resistance in mice (Bohrer et al., 2021). However, it is unclear when granulocytes first respond to Mtb infection of alveolar macrophages with lung infiltration. To quantify the eosinophil response to Mtb infection in the airways of rhesus macaques over time, we used eosinophil-specific intracellular flow cytometric staining of the granule protein eosinophil peroxidase (EPX) as previously described (Bohrer et al., 2021). Using this technique, we quantified eosinophils in bronchioalveolar lavage (BAL) fluid before and after bronchoscopic Mtb-installation and found eosinophils to be significantly increased as early as 7-14 days post infection, at levels ranging from 1%-15% of all CD45 positive immune cells recovered **(Figure 1A)**. To investigate the kinetics of granulocyte recruitment after Mtb infection, and to be able to directly compare eosinophils to neutrophils in the same samples, we devised a NHP granulocyte staining approach based on differential expression of EPX by eosinophils and myeloperoxidase (MPO) by neutrophils within a NHP granulocyte gate **(Figures 1B and S1)**. The NHP granulocyte-gate was based on positive expression of CD11b and CD66abce on granulocytes and exclusion of CD68 expressing monocytes and macrophages **(Figure S1)**. In uninfected rhesus macaque whole blood (WB) eosinophils represented about 10% of all CD11b^+^CD66abce^+^ granulocytes with an eosinophil to neutrophil ratio (E/N) of 0.1. In comparison, eosinophils were relatively enriched in uninfected BAL where they comprised around 40-50% within the granulocyte gate and exhibited a significantly higher E/N ratio (E/N of 1) when compared to blood **(Figure 1B).** To address the possibility that eosinophils were recruited non-specifically in response to micro-tissue damage after bronchoscopy rather than in response to Mtb infection, we performed consecutive bronchoscopies at days -14 and -7 and monitored airway granulocyte influx prior to infection. We found that E/N ratios were largely stable in BAL collected during these initial bronchoscopies, suggesting bronchoscopic lavage by itself was not sufficient to induce rapid changes in airway granulocyte composition **(Figure 1C)**. Notably, eosinophils, and not neutrophils, rapidly increased within seven days of Mtb-instillation and completely dominated the granulocytic response in the airways of rhesus macaques by two weeks, with eosinophils accounting for over 90% of the NHP granulocyte gate in all animals examined **(Figure 1C**). This peak of eosinophils between one to three weeks after exposure was followed by a decline in E/N ratio over the course of two to three months back to or below baseline levels **(Figure 1C**). Our longitudinal data reveals that the initial airway response to Mtb in NHPs is primarily comprised of eosinophilic, and not neutrophilic granulocytes. We next compared the E/N ratio amongst all CD45+ immune cells in BAL to ratios in peripheral WB over time. When eosinophils enriched in the lung airways around two weeks post infection the circulating WB E/N ratios declined at the same time in all animals **(Figure 1D)**. Based on this data we propose a hypothesis whereby circulating eosinophils (either directly or indirectly) sense respiratory Mtb infection and are recruited to the lung and airways from the lung capillary bed, rather than systemic sensing of Mtb infection resulting in increased global eosinophil-output into the blood. Taken together, these findings demonstrated that after Mtb exposure eosinophils and not neutrophils dominate the early granulocytic airway response in rhesus macaques and suggest that eosinophils are rapidly and preferentially recruited from the circulating blood pool to the site of Mtb infection.

**Figure 1:**
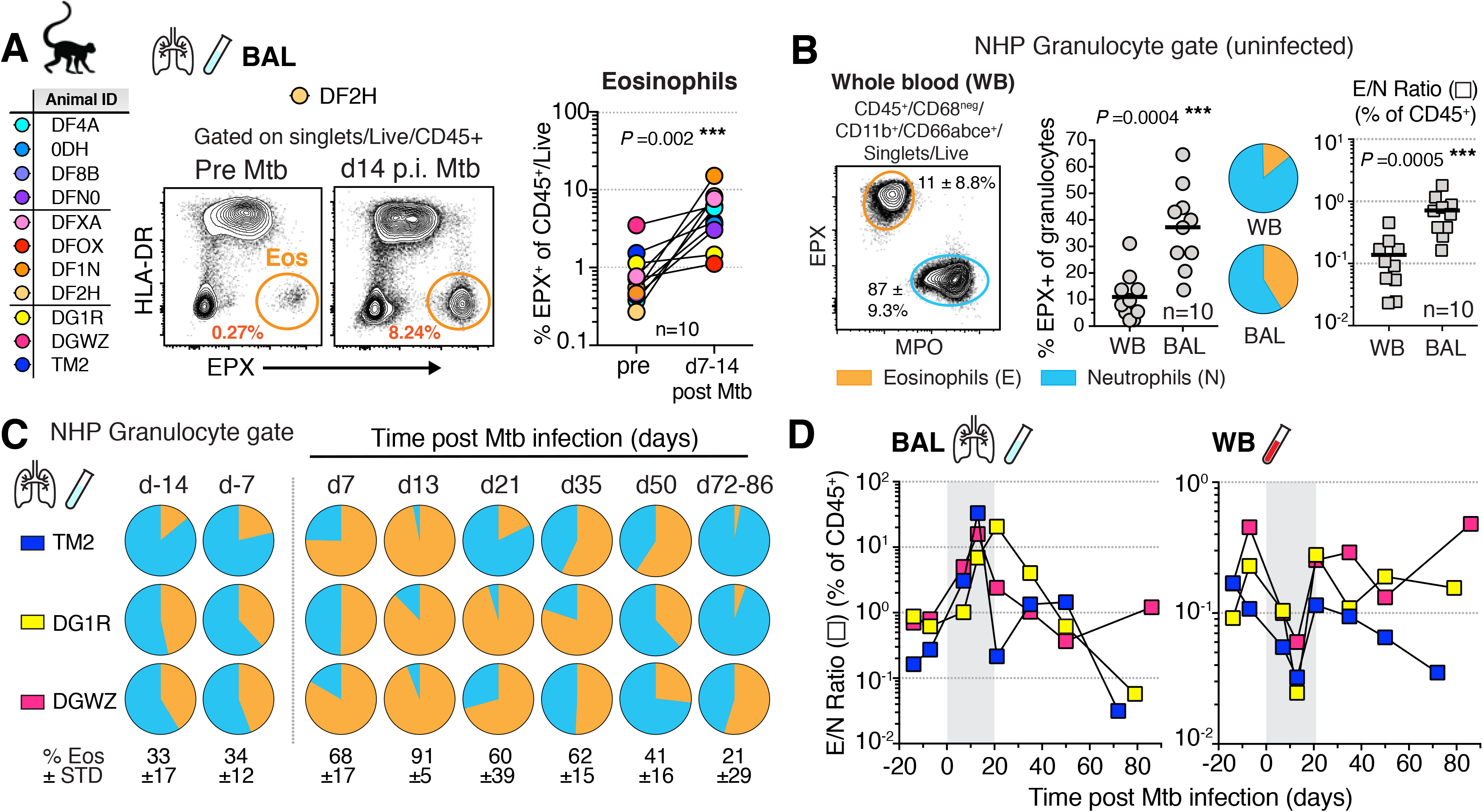
Eosinophils dominate the early granulocytic response in the airways of Mtb-infected rhesus macaques. (**A**) Animal ID list of Mtb infections in rhesus macaques with representative FACS plots of EPX in BAL and quantification of eosinophils before and 1-2 wks. p.i. (n=10, M+F, Matched Wilcoxon) (**B**) NHP granulocyte gating strategy and representative FACS plot with proportion of eosinophils (orange) and neutrophils (blue), alongside eosinophil/neutrophil ratio (E/N) in WB and BAL of uninfected rhesus macaques (n=10, M+F, Matched Wilcoxon) (**C**) NHP Granulocyte gate frequency of eosinophils in BAL prior to and throughout Mtb infection, (n=3, ±SD standard deviation) (**D**) E/N ratio in NHP BAL and WB throughout Mtb infection (n=3, d7-14 time points are highlighted in grey).

### Eosinophils are recruited to Mtb-infected lungs early after infection in mice and interact with Mtb-infected cells in the airways

To provide a model for investigating the underlying mechanisms of eosinophil recruitment we next asked whether eosinophils responded rapdily to low dose aerosol Mtb infection in C57BL6 mice. We assessed granulocyte lung migration by multi-parameter flow cytometry using an intravenous (i.v.) labelling technique which allows the distinction between cells located in the lung vascular capillary bed from cells that trans-migrated into the lung parenchymal tissue or airways (Anderson et al., 2014). Before euthanasia a fluorescently labelled CD45-antibody was injected labeling circulating (i.v.^pos^) cells while lung parenchymal cell populations remained unstained (i.v.^neg^). Eosinophils rapidly accumulated in the lung as early as four days (d4) after Mtb infection **(Figures 2A and S2A)**, a timepoint when Mtb resides almost exclusively in AM (Rothchild et al., 2019), and eosinophils were enriched specifically in the lung vasculature **(Figure 2B)**. Neutrophils numbers or frequencies, in contrast, did not change until two weeks after infection **(Figure 2A)**. Thus, eosinophil numbers and frequencies changed more than a week prior to detectable changes in lung neutrophil numbers and frequencies. Moreover, eosinophils began to migrate into the lung parenchyma around d7-9 post infection (p.i.) with maximum parenchymal eosinophil infiltration at day 14 **(Figures 2B and 2C)**. At this d14 time point, parenchymal neutrophil migration and neutrophil numbers started to dramatically expand with a peak at d21, concurrent with a decline in eosinophil frequencies and numbers **(Figures 2C and S2B)**. Of note, the overall pulmonary E/N ratio in mice **(Figure S2C)** was orders of magnitude lower compared to ratios reported in human lung tissue (Bohrer et al., 2021), NHP TB granulomas (Bohrer et al., 2021) or NHP BAL reported here. So, despite mice exhibiting more neutrophilic lung responses to Mtb infection when compared to humans and rhesus macaques, we still observed an early eosinophil influx. Moreover, this rapid increase in lung eosinophil number and frequency on day 4 without parenchymal migration suggested that eosinophils were sequestered in the capillary bed from circulation. To explore whether the early accumulation of eosinophils was due to local sequestration or instead associated with a systemic increase in peripheral blood we quantified eosinophils in lung and blood. Reminiscent of the dynamics observed in NHP, four days after Mtb infection in mice, eosinophil frequencies decreased in peripheral blood while they simultaneously increased in the lung vasculature bed **(Figure 2D).** Our data in both NHP and mice infected with Mtb thus support the hypothesis that local sequestration, rather than systemic eosinophilia, underlies the concomitant eosinophil increase in the lungs at the earliest stages of Mtb infection. Moreover, our findings suggest a potential sentinel function for circulating eosinophils, that affords rapid detection of bacterial lung infection and eosinophil-retention in the lung vasculature bed and finally migration into the infected tissue.

**Figure 2:**
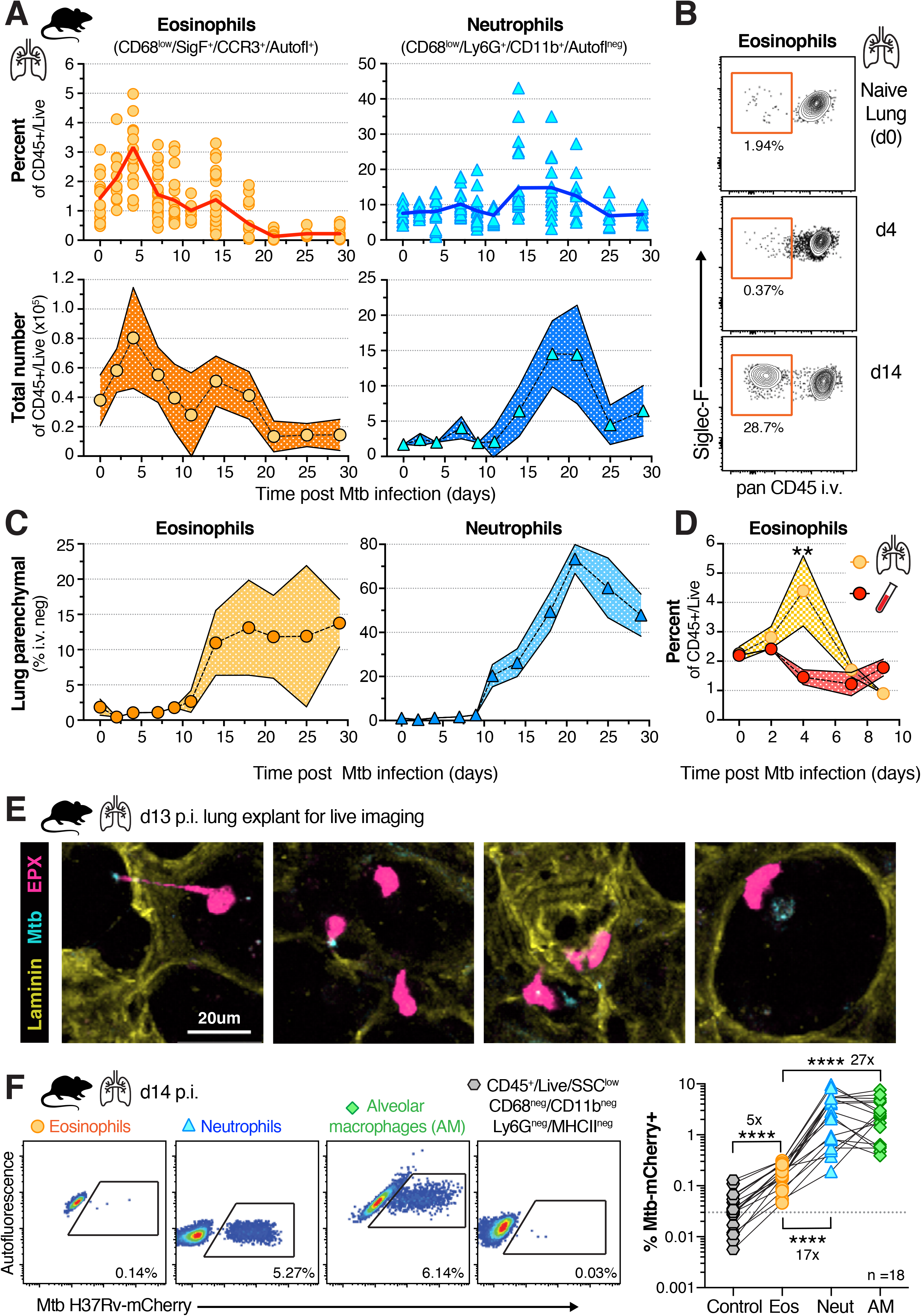
Eosinophils are recruited to lungs early after Mtb infection in mice and interact with Mtb-infected cells in the airways. (**A**) Percent and cell number of total lung eosinophils and neutrophils over time after aerosol infection (100-300 CFU) in WT mice (n=6-25 per time-point, M+F, 95%CI, SEM). (**B**) Representative FACS plot of intravascular staining of eosinophils over time in mouse lung. (**C**) Percent of lung parenchymal (CD45 i.v. neg) eosinophils and neutrophils over time after Mtb infection (n=6-25 per time-point, M+F, 95%CI, SEM). (**D**) Frequency of eosinophils in lung and whole blood after Mtb infection (n=6-10 per time-point, M+F,95%CI, SEM, d4 p=0.007** Mann-Whitney). (**E**) Lung explant model of d13 Mtb-CFP (blue) infected mice expressing an EPX-eosinophil reporter (pink), representative still images (20 μm) from supplemental movie (**F**) Representative FACS plots and quantification of H37Rv-mCherry containing cells in lung on d14 (n=18, M+F) of eosinophils (orange), neutrophils (blue), AM (green) or gated on non-phagocytic, non-myeloid-lineage control cells (grey), dotted line is average signal of control cells (Wilcoxon-matched pairs test for eosinophil comparisons, p<0.0001****).

Since eosinophil migration into the lung parenchyma peaked around d14, we hypothesized that recruited eosinophils could sense or perhaps interact with Mtb or tissue-residing Mtb-infected cells. We used live-imaging of *ex vivo* lung explants from Mtb-Cyan Fluorescent Protein (CFP) infected eosinophil EPX-reporter mice to visualize dynamic interactions between eosinophils and Mtb-containing cells. We observed eosinophils directly interacting with Mtb or Mtb-infected cells in alveoli, moving towards and slowing down next to infected cells as well as extending pseudopods towards bacilli **(Figure 2E, and SM1)**. We then quantified bacteria-containing eosinophils by single cell analysis of fluorescent Mtb-mCherry and found the frequency of Mtb-positive eosinophils (average of 0.15%) to be significantly increased above background assessed in a non-phagocytic SSC^low^, MHCII^neg^, CD68^neg^, CD11b^neg^, Ly6G^neg^ control population (average of 0.03%) of the same sample **(Figure 2F)**. Nevertheless, and in agreement with observations by others (Lee et al., 2020a; Rothchild et al., 2019), two weeks after Mtb infection the proportion of infected neutrophils was 17x and infected AM 27x higher than that of eosinophils within the same sample. Thus, while eosinophils are unlikely to represent a major cellular niche for Mtb or its replication, we show here that they can directly interact with Mtb and Mtb-infected cells at the site of infection.

### CCR3 expression by eosinophils is dispensable for their migration into the lung parenchyma after Mtb infection

Given that Mtb infection is characterized by robust type-I immune responses we next sought to understand the underlying molecular mechanisms leading to lung migration of eosinophils. While eosinophils express a variety of chemotactic receptors, the chemokine receptor CCR3, which binds to eotaxins 1-3, is considered the key regulator mediating eosinophil migration into tissues (Felton et al., 2021; Rothenberg, 1999; Travers and Rothenberg, 2015). So we asked whether type-II associated eotaxins were upregulated after Mtb infection. We initially compared eotaxin mRNA expression in resected human TB lesions from a previously published cohort were we observed a significant increase in eosinophils in metabolically active lesions with high 2-deoxy-2-fluorine-18-fluoro-D-glucose positron emission tomography/computed tomography (^18^FDG PET/CT) uptake compared to low uptake PET lesions in the same individual (Bohrer et al., 2021). We found that eotaxin-2 (CCL24) was significantly reduced in PET high uptake lesions when compared to low uptake PET lesions **(Figure 3A)**. Eotaxin-1 (CCL11), eotaxin-3 (CCL26) and CCR3 itself remained unchanged **(Figure 3A)**. We next confirmed that rhesus macaque eosinophils express CD193 (CCR3) in TB granulomas and BAL **(Figure 3B)** and measured eotaxin levels longitudinally in BAL after Mtb infection **(Figure S3A)**. However, while the E/N ratio significantly positively correlated with eosinophil granule derived EPX and major-basic-protein (MBP) protein levels in BAL across all timepoints **(Figure S3B)**, neither CCL11 nor CCL26 levels associated with the presence of eosinophils **(Figure 3C)**. We then measured eotaxin expression over time after Mtb infection in mice and observed slightly elevated expression of *CCL24* and *CCL26* but not *CCL11* during the first two weeks of infection **(Figure S3C)**, the time frame when eosinophils migrated into Mtb-infected lungs. To functionally test *in vivo* whether chemotactic eotaxin signaling through CCR3 expression on eosinophils was required for early migration into Mtb-infected lungs, we generated competitive mixed bone marrow (BM) chimeras with CCR3-deficient and wild-type (WT) BM **(Figure 3D)**. Competitive mixed BM chimeric mice allow for the side-by-side comparison of the migratory capacity of WT vs. KO eosinophils into the same tissue, thereby normalizing bacterial burden and inflammatory milieu which likely impact cellular migration during Mtb infection. Two weeks after infection we assessed the ability of WT or CCR3-deficient eosinophils to transmigrate into the lung parenchyma and observed significantly higher frequencies of parenchymal *Ccr3^-/-^* eosinophils compared to WT eosinophils in the same animals **(Figure 3E)**. When CCR3 expression on eosinophils was absent, their migration into lung tissue increased more than two-fold, arguing that CCR3-dependent signals in fact limited rather than promoted migration into the Mtb-infected lung **(Figure 3F)**. Thus, despite detectable expression of type-II chemotactic ligands, early eosinophil migration into Mtb-infected lung tissue in mice does not require classical CCR3-dependent chemotaxis and eosinophil abundance did not correlate with eotaxin levels in the BAL of Mtb infected rhesus macaques.

**Figure 3:**
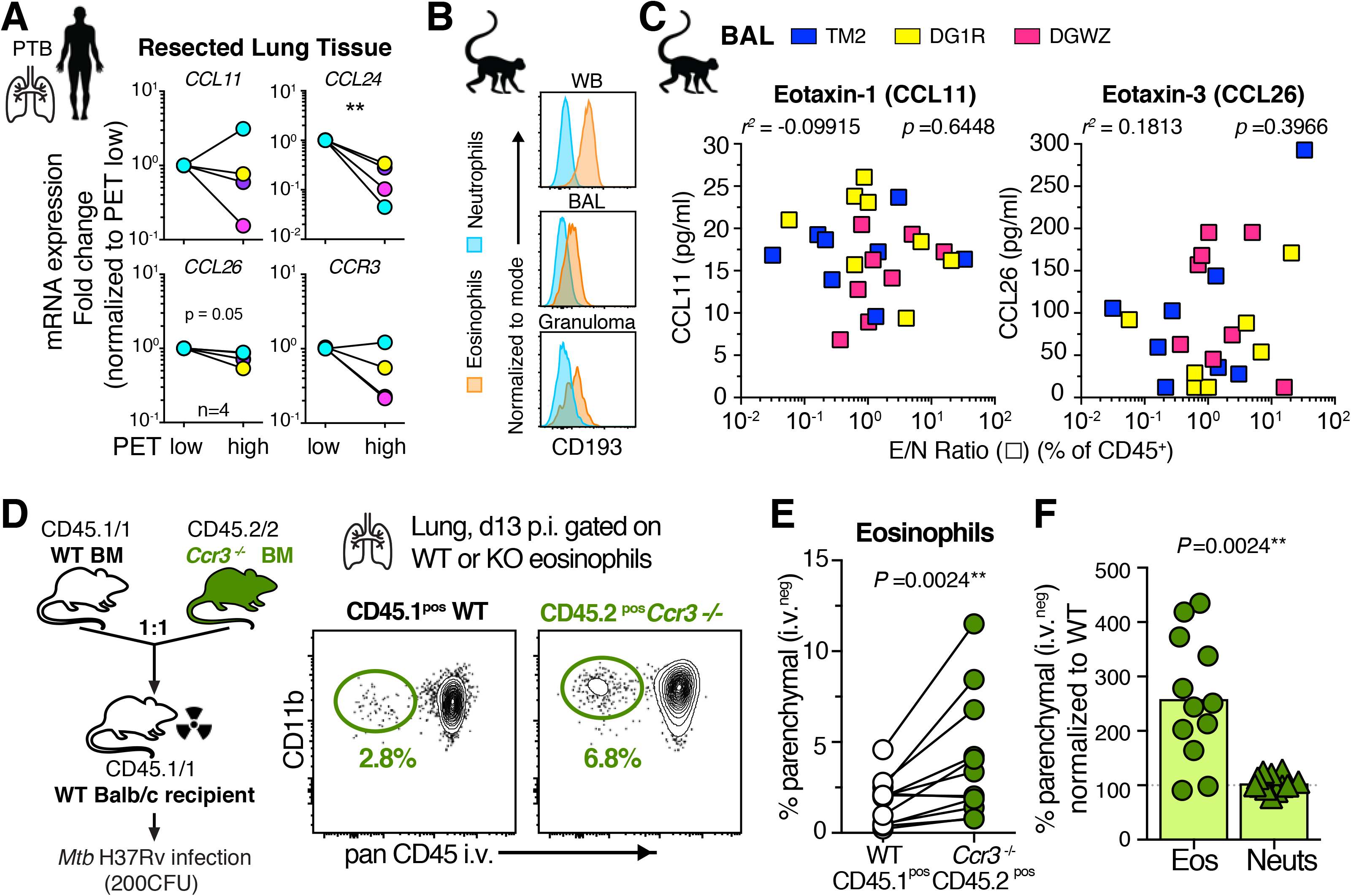
CCR3 expression on eosinophils is dispensable for their migration into the lung parenchyma after Mtb infection. (**A**) qRT-PCR of indicated genes in ^18^FDG PET/CT low or high signal intense human TB lung lesions (n=4, Wilcoxon-matched pairs, **p<0.01) (**B**) Example CD193/CCR3 expression of granulocytes after Mtb infection in rhesus macaques (**C**) Eotaxin levels in BAL of Mtb infected rhesus macaques across all timepoints correlated with E/N ratio in same samples (Spearman). (**D**) Balb/c *Ccr3*^-/-^ competitive mixed BM chimeric mice (mBM) and representative FACS plot of lung parenchymal eosinophils after Mtb infection (**E**) Quantification of CD45 i.v. negative granulocytes in mBM lung (M+F, n=12, Wilcoxon-matched pairs) (**F**) Migration efficiency of granulocytes normalized to WT cells in same lung (Wilcoxon-matched pairs).

### Eosinophils express high levels of the oxysterol sensor GPR183 and GPR183 expression is significantly reduced in peripheral blood during active TB in patients

Granulocytes can migrate in response to a variety of G-protein-coupled receptor (GPR) ligands such as chemokines, complement components and lipid mediators (Rosenberg et al., 2013; Shamri et al., 2011). GPR183, also known as Epstein-Barr virus-induced GPR 2 (Ebi2), is a chemotactic receptor expressed on T and B lymphocytes, macrophages, DCs, astrocytes and innate lymphoid cells and has been most extensively studied in cell positioning in lymphoid organs (Barington et al., 2018; Cyster et al., 2014; Hannedouche et al., 2011; Li et al., 2016; Liu et al., 2011; Pereira et al., 2009; Rutkowska et al., 2016; Willinger, 2019; Yi et al., 2012). GPR183 recognizes and binds to oxysterols which are oxidized versions of cholesterol or its molecular precursors that can have biological and chemotactic activity (Cyster et al., 2014; Schroepfer, 2000). Consistent with western blot detection of GPR183 in human eosinophils (Shen et al., 2017), we report here uniformly high levels of GPR183 expression on eosinophils at the single cell level by flow cytometry in peripheral blood of rhesus macaques **(Figure 4A)** and healthy individuals **(Figure 4B)**. After Mtb infection in macaques GPR183 expression levels remained consistently high on eosinophils and negative on neutrophils in TB granulomas **(Figure 4C)**. To assess GPR183 expression on lung eosinophils in patients, we measured GPR183 levels by flow cytometry in freshly isolated eosinophils from resected TB lung lesions of pulmonary TB (PTB) patients **(Figure S4A)**. Again, eosinophils displayed the highest GPR183 expression compared to macrophages, T or B lymphocytes and GPR183 expression did not change with ^18^FDG PET/CT signal intensity **(Figure 4D)**. Cholesterol-25-hydroxylase (Ch25h) is a key enzyme involved in the generation of oxysterol ligands for GPR183 and both global *CH25H* and *GPR183* mRNA expression levels in lung tissue were unchanged in ^18^FDG PET/CT high compared to low signal intensity lesions **(Figure S4B)**. Of note, when we explored transcriptome profiles of 961 blood samples with 433 from latent TB infection and 528 active TB across eight independent published cohorts, *GPR183* was significantly downregulated in active PTB patients compared to latent TB controls **(Figure 4E)**. Further, in an adolescent cohort followed longitudinally during disease progression (Zak et al., 2016) significant downregulation of *GPR183* preceded diagnosis of active TB by six months **(Figure 4F)**. These data suggest that GPR183 expression itself, and/or GPR183-expressing circulating leukocytes are modulated in the blood during progression to active TB disease in patients.

**Figure 4:**
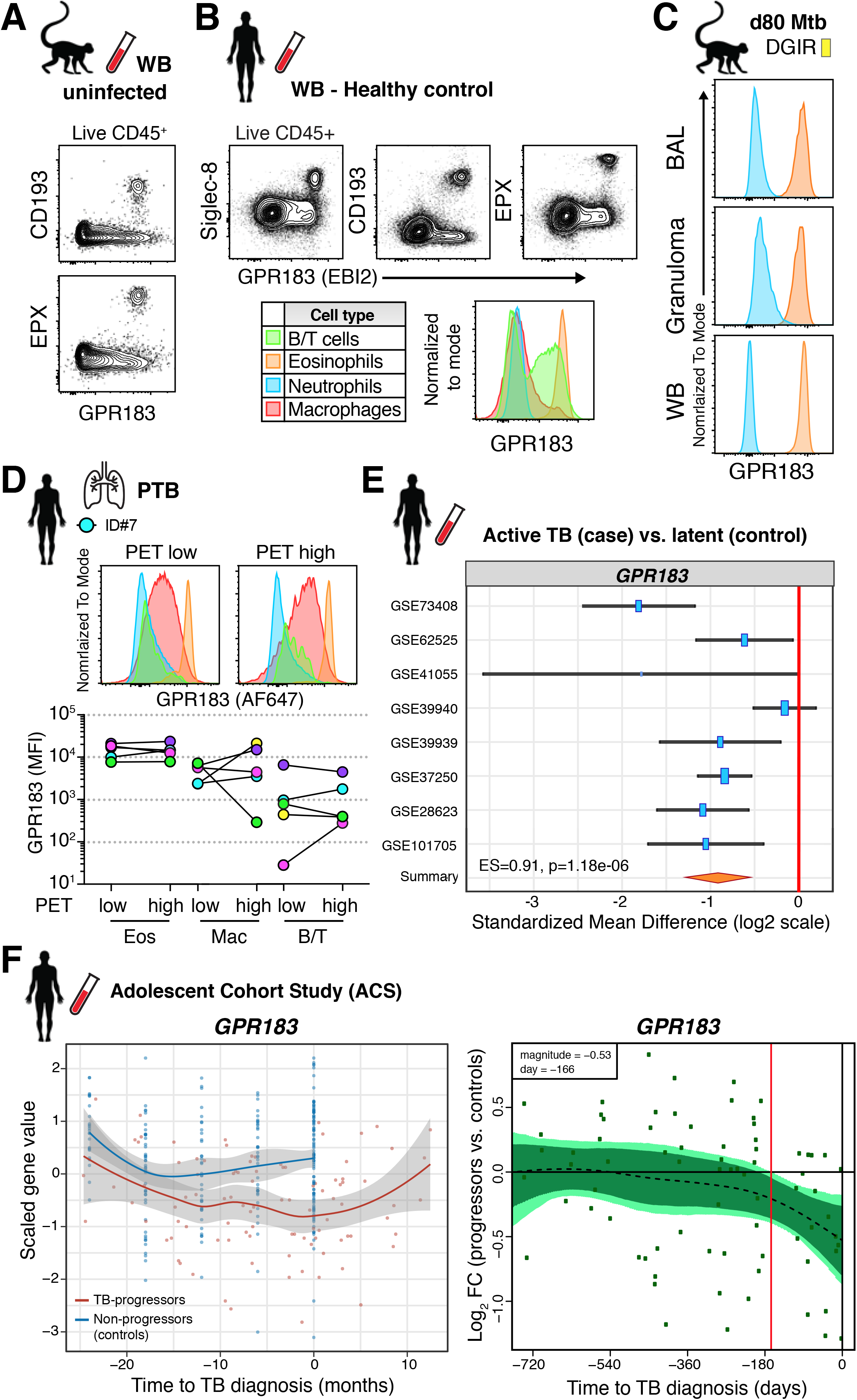
Human and rhesus macaque eosinophils express uniformly high levels of the oxysterol receptor GPR183. (**A**) Representative FACS plots of GPR183 expression on eosinophils in WB of uninfected macaques (**B**) Representative FACS plots of GPR183 expression on leukocytes and eosinophils in WB from healthy donors (**C**) Example GPR183 expression of granulocytes after Mtb infection in BAL, WB and granuloma of macaque DG1R (**D**) GPR183 expression by flow cytometry on immune cells in ^18^FDG PET/CT low or high FDG uptake in human TB lung lesions (**E**) *GPR183* expression from transcriptome profiles of LTBI controls (n=433) or ATB (n=528) blood samples across 8 indicated cohorts (z-test on summary effect size computed as Hedges’ g). (**F***) GPR183* expression kinetics over time from Adolescent cohort study. Left side graphs depicts treatment follow-up from 42 subjects that progressed to active TB (red) and 109 latently infected controls (blue), (local regression model, Hedge’s g) and *GPR183* mRNA expression over time, expressed as log2 fold change (FC) between bin-matched progressors (n = 44) and controls (n = 106) and modeled as nonlinear splines (dotted line). Right side graph shows progressors with light green shading representing 99% CI and dark green shading 95%CI temporal trends, computed by performing 2000 spline fitting iterations after bootstrap resampling from the full data set. Deviation time (day -166) is calculated as the time point at which the 99% CI deviates from a log2 fold change of 0, is indicated by the vertical red line.

### Oxysterol ligands are dynamically regulated in the airways of Mtb-infected rhesus macaques and can mediate GPR183 dependent eosinophil migration *in vitro*

GPR183 is known to permissively bind oxysterols hydroxylated at the 7, 25, and 27 position with a preference for 7α,25-dihydroxycholesterol (7α,25-di-OHC) (Hannedouche et al., 2011; Liu et al., 2011). To ask whether Mtb infection resulted in changes in oxysterol metabolism we directly measured oxysterols in the BAL before and after Mtb infection in rhesus macaques by liquid chromatography tandem mass spectrometry (LC-MS/MS) using previously described methodology (McDonald et al., 2012). Unesterified oxysterol signals that were present in >50% of samples were included for analysis. Consistent with its reported dissociation constant in the high picomolar range and accounting for BAL-associated dilution of the alveolar fluid, 7α,25-di-OHC measurements were below our ∼5 nM limit of detection. In two of the three examined animals, hydroxycholesterol levels exhibited time-dependent dynamics post infection with changes in some classes evident as early at 7 days post-infection **(Figure 5A)**. While we found no association of eotaxins with E/N ratios across all time points, two oxysterols significantly correlated with the E/N ratio in Mtb-infected NHP BAL. 7α,27-di-OHC, a GPR183 ligand with a mid-nanomolar dissociation constant correlated negatively and an oxidized derivative of 7α-OHC, 3-oxo-7α-hydroxycholesterol correlated positively with E/N ratios **(Figure 5B).** The biological functions of the various oxysterol metabolites *in vivo* are not well understood, and optimization of oxysterol airway sampling and detection are needed to fully elucidate GPR183 signaling topology at the ligand level *in vivo*. Nevertheless, the increase of 3-oxo-7α-hydroxycholesterol may be indicative of more broadly 7α-substituted cholesterol metabolism with increasing E/N ratios. One possible interpretation of the negative correlation of 7α,27-di-OHC with relative eosinophil abundance could be the occurrence of ligand-depletion as the influx of target cells increases. To confirm that oxysterol ligands could directly trigger GPR183-dependent migration of eosinophils we used an *in vitro* migration assay (Hong et al., 2020; Shen et al., 2017). We detected significant migration when purified human eosinophils were exposed to 7α,25-di-OHC but not 25-hydroxycholesterol (25-OHC) *in vitro*, and this response was abrogated with addition of the GPR183-specific inhibitor NIBR189 **(Figure 5C)**. These data demonstrate that GPR183 engagement on eosinophils promotes chemotaxis and thus may contribute to cellular recruitment of eosinophils to the lungs during Mtb infection.

**Figure 5:**
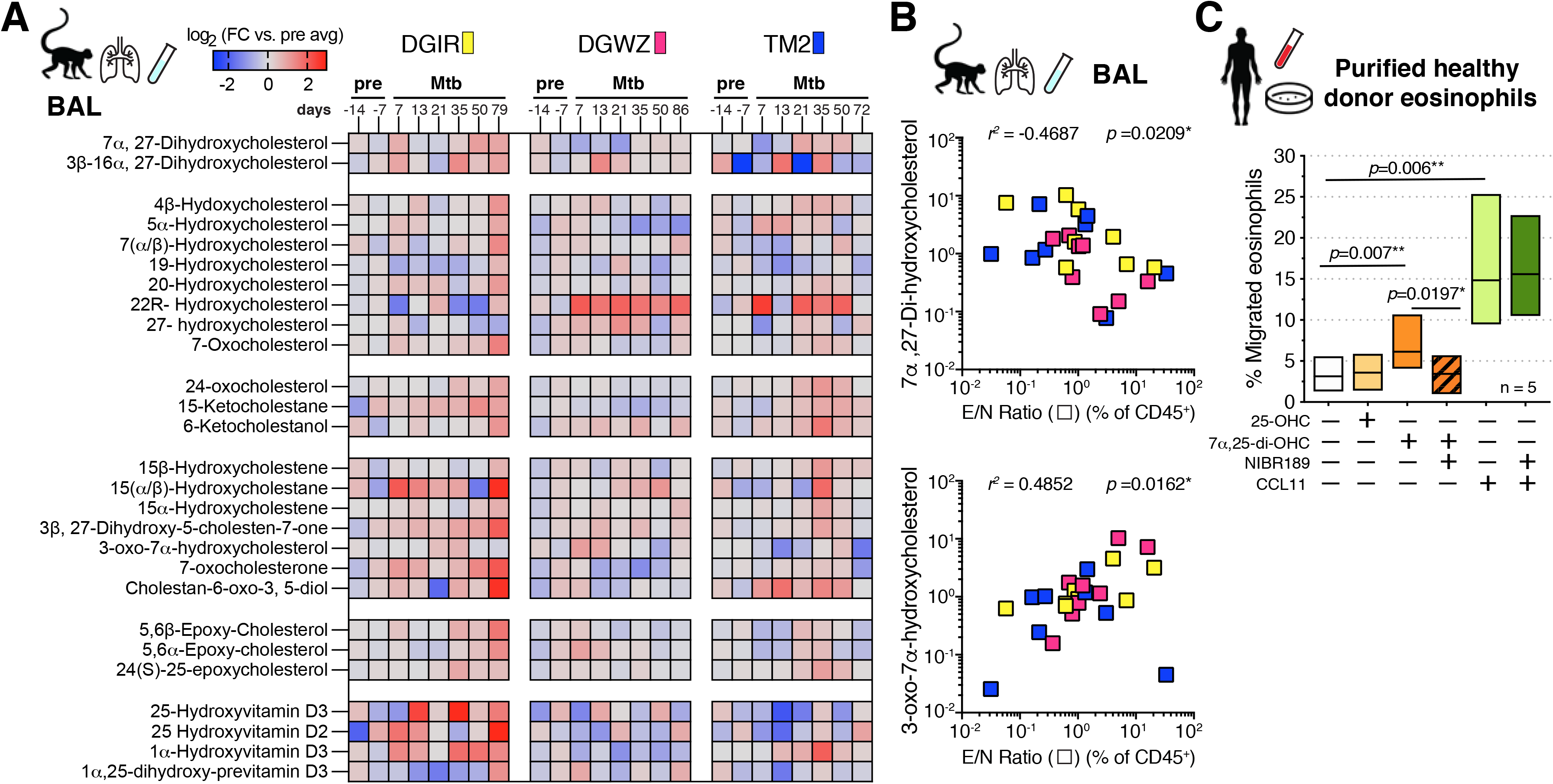
Oxysterol GPR183-ligands are dynamically regulated in rhesus macaque airways after Mtb infection and trigger migration of eosinophils *in vitro*. **(A)** Oxysterol metabolite signals in BAL of Mtb-infected rhesus macaques over time displayed as log_2_ fold change vs. average pre-infection time point-intensity measured by LC-MS/MS (n=3), **(B)** 7α 27-di-OHC and 3-oxo-7α-OHC levels in BAL of Mtb-infected rhesus macaques across all timepoints correlated with E/N ratio in same samples (Spearman). **(C)** *In vitro* migration assay of purified human eosinophils to oxysterol ligands 25-OHC (100nM) and 7α CCL11 (eotaxin-1, 100ng/ml) in the presence or absence of NIBR189 (50nM), a selective GPR183 inhibitor (n=5, Wilcoxon-matched pairs)

### Ch25h derived oxysterols mediate eosinophil-intrinsic GPR183 dependent migration into Mtb-infected lungs *in vivo*

To directly test whether oxysterols mediate lung migration of eosinophils *in vivo*, we initially asked if *Ch25h* mRNA is induced after Mtb infection in mice. Whole lung *Ch25h* mRNA expression levels modestly increased in the second week of Mtb exposure **(Figure 6A**), a time point when we saw peak eosinophil migration into lungs. However, we observed eosinophil sequestration in the lung vasculature bed as early as 4 days after low dose aerosol infection of mice. At this very early stage Mtb resides almost exclusively in AM and we thus hypothesized that infected AM may upregulate Ch25h to alert circulating eosinophils of Mtb infection. In fact, in a recently published RNASeq dataset in a high dose Mtb infection model Mtb-infected vs. uninfected AM *Ch25h* expression was significantly increased in infected cells as early as day four post infection **(Figure 6B)** (Rothchild et al., 2019). Upregulation of other enzymes involved in oxysterol biosynthesis (*Cyp7a1*, *Cyp7b1, Cyp8b1, Cyp3a11*) or the eotaxins *Ccl11*, *Ccl24* or *Ccl26* was not detected in infected AM on day 4 or 10 post infection in the same dataset (Rothchild et al., 2019). To test the hypothesis that Ch25h-dependent oxysterols regulate pulmonary eosinophil localization as early as 4 days after low dose Mtb infection *in vivo*, we quantified the observed early enrichment of eosinophils in the lung vasculature **(Figures 2A-C)** in *Ch25h^-/-^* and *Gpr183^-/-^* mice. Indeed, the earliest eosinophil accumulation was significantly reduced in both *Ch25h^-/-^* and *Gpr183^-/-^* mice **(Figure 6C)**. We then asked whether Mtb-harboring AM selectively upregulate *Ch25h* expression at two weeks after low dose Mtb aerosol infection, the time point where we observed eosinophil interactions with Mtb-infected cells by live imaging and maximum parenchymal eosinophil migration. To this end we FACS purified Mtb-infected and uninfected AM from BAL pools after low dose aerosol infection and measured relative *Ch25h* expression by qPCR **(Figure 6D)**. We found that infected AM expressed significantly higher levels of *Ch25h* than uninfected bystander AM from the same BAL samples **(Figure 6D)**.

**Figure 6:**
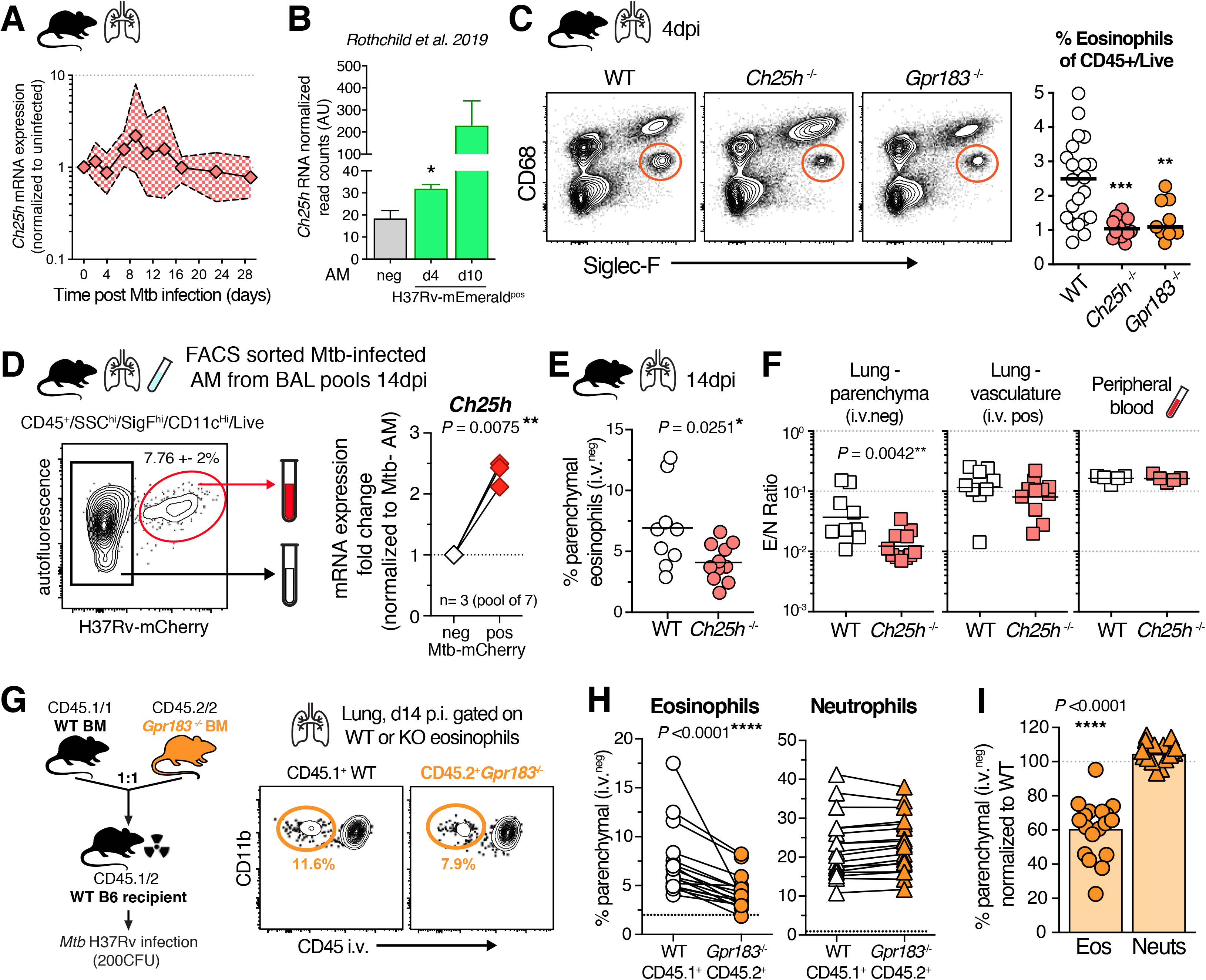
Ch25h derived oxysterols mediate eosinophil-intrinsic GPR183-dependent extravasation into Mtb-infected lungs *in vivo*. (**A**) qRT-PCR *Ch25h* expression in lung tissue after aerosol Mtb infection in WT mice (n=5-12). (**B**) *Ch25h* mRNA read arbitrary units (AU) based on data published by Rothchild et al. 2019, normalized to uninfected bystander reads, after H37Rv-mEmerald high dose (2000-4000 CFU) Mtb infection in mice (two-tailed unpaired t-Test) (**C**) Representative FACS plots of eosinophils in the lungs of *Gpr183*^-/-^, *Ch25h*^-/-^ or WT B6 mice at 4 dpi and quantification (M+F, n=10-20, **p<0.01, ***p<0.001, Mann-Whitney) (**D**) Representative FACS plots of low dose aerosol Mtb-infected (100-300 CFU) WT B6 AM at 14dpi from BAL and *Ch25h* gene expression in purified AM populations (M+F, n=3 x pool of 7, Wilcoxon-matched pairs) (**E**) Percent lung parenchymal eosinophils d14pi in *Ch25h*^-/-^ mice (M+F, n=9-10, Mann-Whitney) (**F**) E/N ratio in lung parenchymal and lung vascular CD45+ immune cells compared to the peripheral blood at 14dpi in *Ch25h*^-/-^ mice (M+F, n=9-10, Mann-Whitney) (**G**) WT and *Gpr183*^-/-^ competitive mixed BM chimeric mice and representative FACS at 14dpi. Mtb infection (**H**) Quantification of CD45 i.v. negative granulocytes in mBM lung (M+F, n=18, Wilcoxon-matched pairs, dotted black line shows average CD45 i.v. negative eosinophils or neutrophils in uninfected mixed BM chimeric mice).

These data support the hypothesis that Mtb itself triggers the upregulation of the oxysterol producing enzyme Ch25h in infected AM. When we assessed eosinophil extravasation into the lung parenchyma two weeks after Mtb infection, the frequency of parenchymal eosinophils was significantly reduced in *Ch25h*^-/-^ mice compared to WT animals **(Figure 6E)**. The decrease in tissue residing eosinophils was reflected in selectively and significantly reduced E/N ratios in the lung parenchyma but not the lung vasculature or peripheral blood **(Figure 6F)**. Finally, to functionally demonstrate that direct cell-intrinsic oxysterol signaling via GPR183 expression on eosinophils is required for migration into Mtb-infected lungs *in vivo*, we used competitive mixed *Gpr183^-/-^* and WT BM chimeras **(Figures 6G and S5A)**. We consistently observed significantly lower frequencies of parenchymal *Gpr183^-/-^* eosinophils compared to WT eosinophils in each of the 18 animals examined, while GPR183 deficiency failed to alter the migratory capacity of neutrophils **(Figure 6H)**. *Gpr183*^-/-^ eosinophils exhibited on average a 40% reduction in their tissue migration compared to WT *Gpr183^+/+^* eosinophils in the same animal **(Figure 6I).** Our data thus show that eosinophils, through cell-intrinsic GPR183 engagement, respond to Mtb infection by early migration into the lung tissue where they can interact with Mtb-infected cells. Lastly, to address whether GPR183 is generally required for optimal lung migration by eosinophils, we assessed the migration capacity of *Gpr183^-/-^* deficient eosinophils in competitive mixed *Gpr183^-/-^* and WT BM chimeric mice in models of pulmonary viral infection and in a type-II inflammation-driven lung allergy. When mixed BM chimeric mice were infected with the B1.351 variant of SARS-CoV-2, *Gpr183^-/-^* eosinophils migrated as efficiently as WT eosinophils into the infected lung 3-5 days post exposure **(Figure S5B)**. Likewise, eosinophil-intrinsic expression of GPR183 was dispensable for migration into the parenchyma in the Ovalbumin (OVA)-Alum lung allergy model **(Figure S5C)**. Taken together our data demonstrate that after Mtb infection the oxysterol generating enzyme Ch25h promotes rapid lung migration of eosinophils through eosinophil-intrinsic expression of the oxysterol receptor GPR183. Our data suggest that oxysterols may play unique roles in both alerting circulating eosinophils to the presence of Mtb and facilitating eosinophil-macrophage interactions.

## DISCUSSION

Intracellular bacterial infections are typically not associated with type-II cell infiltration, yet we recently showed that at later stages of Mtb infection eosinophils populate TB granulomas and contribute to host resistance in mice after Mtb infection (Bohrer et al., 2021). Here, we explored the early lung recruitment dynamics of granulocytes after Mtb infection in mice and rhesus macaques and demonstrate that eosinophils, not neutrophils, are among, if not the first, leukocytes from circulation to respond rapidly to Mtb infection. Not only do eosinophils increase in the lungs within one to two weeks after infection, they also interact with tissue resident Mtb-infected cells based on live imaging of lung explants. We suggest that infected-cell triggered chemotactic signals sound the alarm of an ongoing Mtb infection to circulating eosinophils which then migrate to the infection site. Moreover, we provide a molecular mechanism of eosinophil migration in response to Mtb infection by showing that both Ch25h and eosinophil-intrinsic GPR183 expression mediate their accumulation in the pulmonary vasculature and migration into lung parenchyma. In contrast, GPR183 had no role in eosinophil migration after SARS-CoV-2 infection and in the OVA-alum lung allergy model, supporting the hypothesis that the Ch25h-GPR183 axis may be preferentially utilized by bacteria harboring cells.

Cell-cell interactions of eosinophils with tissue resident macrophages have been described previously in the dermis and adipose where they contribute to tissue homeostasis and maintenance of resident macrophages during infection (Lee et al., 2020b; Shah et al., 2020; Wu et al., 2011). These interactions are often cooperative with macrophages secreting eotaxin to recruit eosinophils which in turn provide IL-4 or IL-13 to macrophages resulting in metabolic reprogramming and cellular maintenance (Lee et al., 2020b; Minutti et al., 2017; Shah et al., 2020; Weller and Spencer, 2017; Wu et al., 2011). Because 1) alveolar macrophages are the initial cellular reservoir of Mtb in the lung (Cohen et al., 2018; Rothchild et al., 2019), 2) we show here that bacilli-harboring AM’s upregulate *Ch25h* expression, 3) *ex vivo* imaging studies observed eosinophils in alveoli interacting with Mtb-infected cells and 4) we found no role for CCR3 in eosinophil recruitment after Mtb infection, we speculate that oxysterols, and not eotaxins, derived from Mtb-infected AM may play a role in the earliest recruitment of eosinophils after infection with Mtb. Nevertheless, our data do not exclude a role for oxysterol production by additional cell types. *Ch25h* expression can be induced by a variety of signals, including TLR stimulation, type I interferons and IL-1 family cytokines (Ahsan et al., 2018; Magoro et al., 2019; Park and Scott, 2010) and Ch25h derived oxysterols produced by pulmonary vascular endothelium, basal epithelium or fibroblasts may also contribute to oxysterol generation (Montoro et al., 2018; Willinger, 2019). The exact cellular compartments in the lung responsible for the Ch25h-dependent oxysterol response and the pathways resulting in *Ch25h* up-regulation of Mtb infected tissue resident macrophages after Mtb infection *in vivo* warrant further study. Moreover, eosinophil-macrophage interactions during Mtb infection are likely bi-directional as well. It is conceivable that eosinophils, which are loaded with diverse preformed cytokines and bioactive lipid mediators, can provide either inhibitory or activating signals to Mtb-infected macrophages in the airways. Accordingly, one plausible signal to explore is eosinophil-derived IL-4 or IL-13. Indeed, macrophage-epithelial transition and TB granuloma formation require IL-4Ra induced STAT6 activation (Cronan et al., 2021). Thus, future studies should aim to identify the relevant cellular interaction partners of eosinophils during early and late stages of Mtb infection and dissect the molecular nature and importance for host resistance.

While we demonstrated that after Mtb infection early eosinophil migration into the lungs is independent of CCR3, additional mechanisms of eosinophil recruitment likely also play a role as *Gpr183*^-/-^ eosinophils still displayed some residual lung-homing ability. Although neutrophils did not utilize GPR183 for lung homing and CD4 T cells have previously been shown to not require GPR183 (Hoft et al., 2019) we cannot exclude a role for GPR183 in the migration of other cell types. In this context, we show that global blood transcriptional expression of *GPR183* is downregulated prior to diagnosis and during active TB disease in patients. Several non-mutually exclusive hypotheses could explain *GPR183* reduction in blood during TB: 1) *GPR183* is downregulated on circulating immune cells, including eosinophils, NK cells, monocytes, T and B cells; 2) GPR183-expressing cells are selectively depleted in circulation; and 3) GPR183-expressing cells leave the circulation to enter inflamed infected tissue sites, by means of either oxysterol or other chemotactic cues. We also cannot exclude a role for GPR183 in cell positioning once cells have arrived in the lungs. For example, GPR183 expression by T and B cells has been extensively studied in immune cell positioning in secondary lymphoid organs (Barington et al., 2018). Thus, akin to oxysterol positioning of B and T lymphocytes in lymphoid follicles, GPR183 could also play a role in aggregating and guiding both adaptive and innate immune cells to Mtb-infected macrophages at later stages inside lung tissue and granulomas.

Lastly, in addition to chemotaxis GPR183 ligands can also exhibit direct antiviral properties as well as amplify inflammatory transcriptional responses in macrophages (Gold et al., 2014; Liu et al., 2013), indicating that GPR183 and Ch25h may contribute to host resistance against Mtb independently of chemotactic functions. However, since both pro and anti-bacterial effects of oxysterol signaling on Mtb-infected macrophages have been described *in vitro* the non-chemotactic functions of oxysterols and their relevance in host resistance to Mtb infection are not yet clear (Ahsan et al., 2018; Bartlett et al., 2020; Tang et al., 2020).

Taken together, our findings provide new mechanistic insight into the early *in vivo* immune response to Mtb-infected alveolar macrophages and reveal an unexpected involvement of eosinophils in both mice and rhesus macaques. Importantly we show that Mtb-infected cells upregulate the oxysterol-producing enzyme Ch25h and that the oxysterol receptor GPR183 expressed on circulating eosinophils triggers lung migration early after pulmonary Mtb infection. Moreover, the data presented here show that in addition to contributing to host resistance in mice at later stages of infection (Bohrer et al., 2021) eosinophils may serve as early first-responder cells during initial Mtb infection.

## ACKNOWLEDGEMENTS

The authors thank the clinical study participants and medical staff at SPHCC affiliated with Fudan University in Shanghai. We thank Drs. David Sacks and Yasmine Belkaid for discussion and feedback on the manuscript and Ryan Kissinger for graphical abstract assistance. We are especially grateful to the staff of the NIAID ABSL2 and ABSL3 facilities, Theresa Hawley, Melanie Cohen and Tom Moyer for FACS based cell sorting under BSL3 conditions and Reed Johnson and the NIAID SARS-CoV-2 Virology Core for viral stocks. This work was supported in part by the intramural research program of NIAID (K.D.M-B, C.M.B, D.L.B, A.S., L.E.V., A.D.K.) and the National Natural Science Foundation of China grant (#81770010 to K-W.W.)

## AUTHOR CONTRIBUTIONS

Conceptualization: A.C.B, E.C., A.D.K, C.M.B., K.D.M.-B.; Methodology: A.C.B, E.C., B.S., M.A.M., F.L., P.J.B, Z.H., L.W.,TBIP, O.K., Y.S., K.-W.W. and K.D.M.-B; Investigation: A.C.B., E.C., C.E.T, M.A., M.A.M., E. B. F.L., K.L.H, P.J.B., F.T-J., Z.H., H.M., L.W., L.N., W.Z., K.D.K., TBIP, O.K., M.D., K.-W.W. and K.D.M.-B.; Resources: TBIP, C.M.B., P.K., T.J.S., A.S., L.E.V., D.L.B., S.H.L., Y.S., A.D.K., K.-W.W. and K.D.M.-B, Data analysis and curation: A.C.B., E.C., M.A.M., F.L., M.D., B.S., C.M.B, T.J.S., P.K., and K.D.M.-B, Writing-original draft: K.D.M.-B, Writing-review and editing: all co-authors, Visualization: A.C.B., E.C., B.S., M.D., T.J.S., P.K. and K.D.M.-B; Supervision: A.D.K, P.K., D.L.B., L.E.V., Y.S., K.-W.W., C.M.B and K.D.M.-B. Funding acquisition: A.S., C.M.B., A.D.K., D.L.B., L.E.V., K.-W.W. and K.D.M.-B.

## SUPPLEMENTAL FIGURE LEGENDS

**Supplemental Figure S1:**
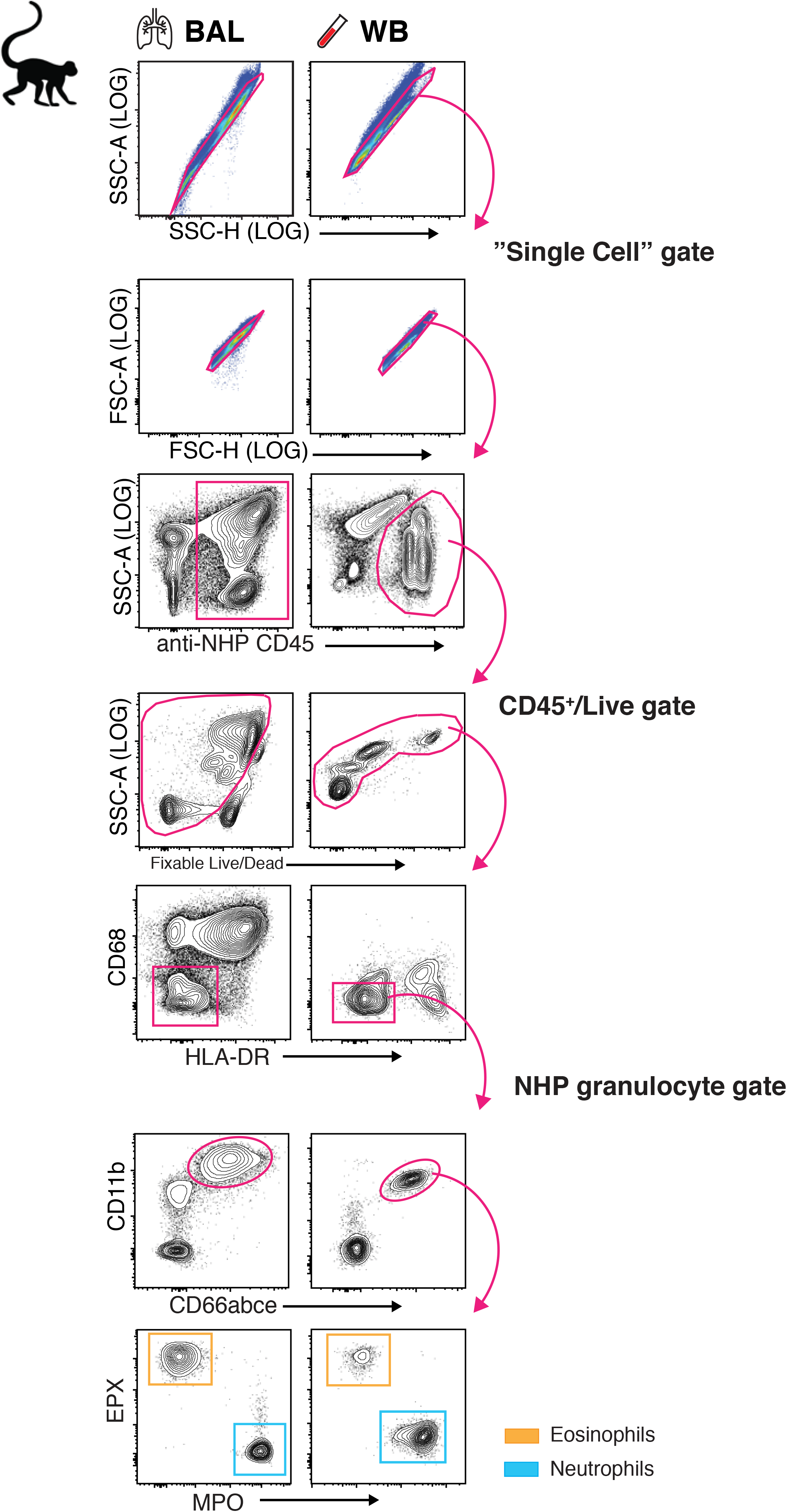
Eosinophil and neutrophil staining and gating strategy in rhesus macaques. (**A**) Representative FACS plots and NHP granulocyte gating strategy in BAL and WB for rhesus macaque eosinophils (orange gate) and neutrophils (blue gate).

**Supplemental Figure S2:**
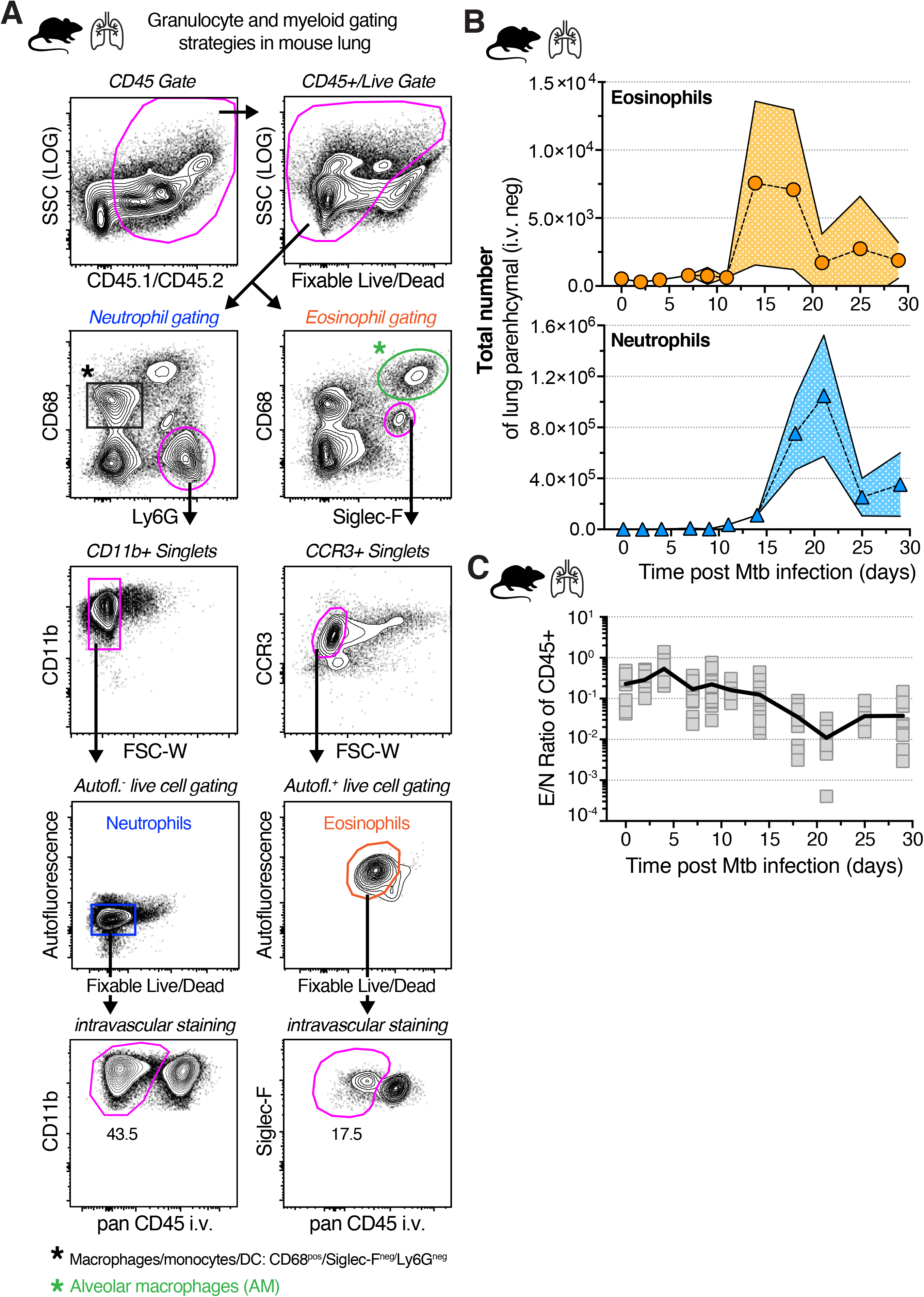
Granulocyte and myeloid cell gating strategy and kinetics of granulocytes after Mtb infection in mice. (**A**) Representative FACS plots and gating strategies in WT B6 mouse lung 14 days after aerosol (100-300 CFU) infection in WT B6 mice for eosinophils (orange), neutrophils (blue), AM (green) and pan-myeloid monocytes/macrophages/DC (black) (**B**) Total cell number of lung parenchymal (CD45 i.v. negative) eosinophils and neutrophils over time after aerosol infection in WT mice (n=6-25 per time-point, M+F, 95%CI, SEM) (**C**) E/N ratio in lung over course of Mtb infection in WT mice (n=6-25 per time-point).

**Supplemental Figure S3:**
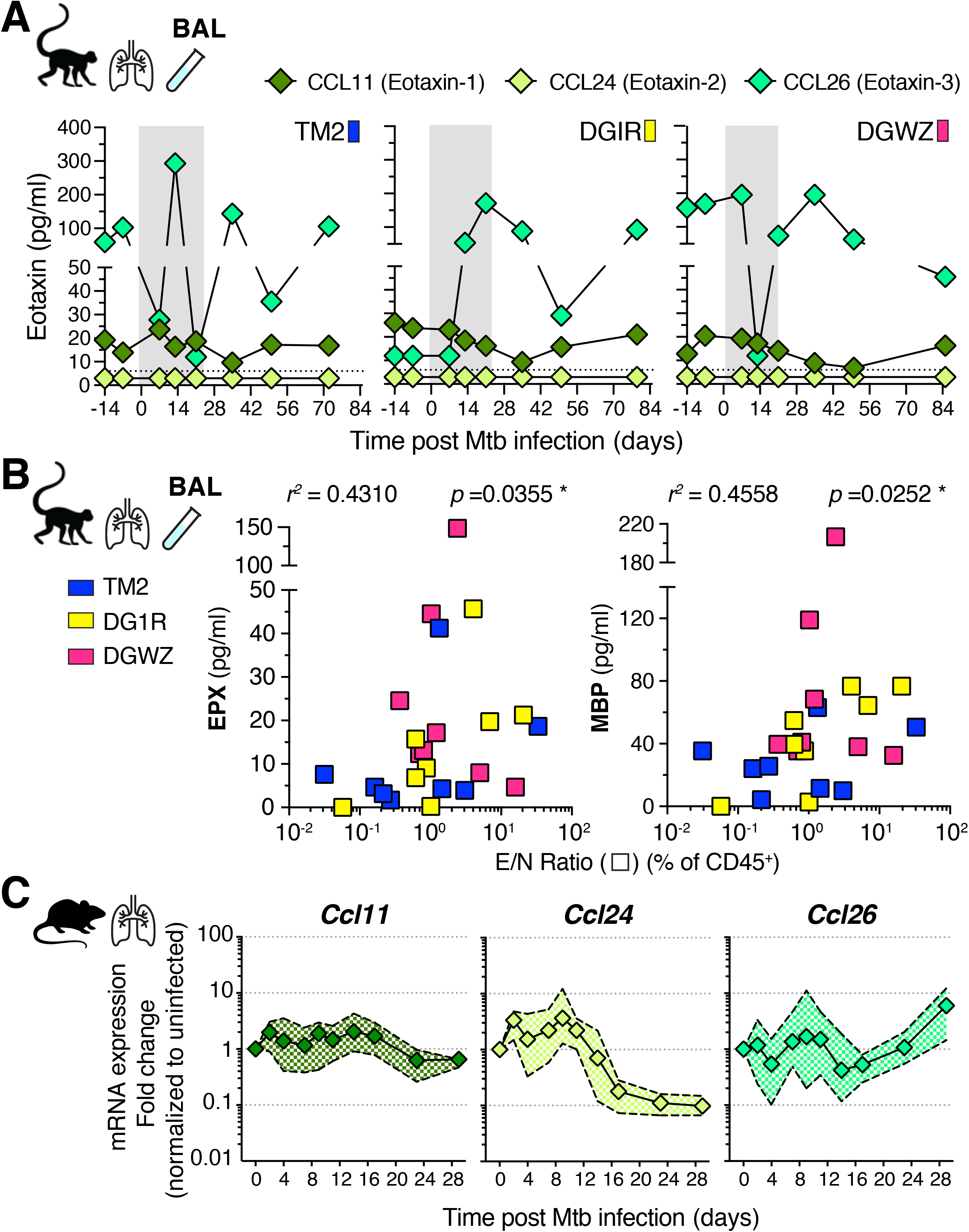
Eotaxin levels in airways and lungs of Mtb infected rhesus macaques and mice. (**A**) Eotaxin levels in BAL of rhesus macaques over the course of Mtb infection, dotted line represents limit of detection, Eotaxin-2 (CCL24) was undetectable (n=3, d7-14 time points are highlighted in grey) (**B**) EPX and MBP (major-basic-protein) levels in BAL of Mtb infected rhesus macaques (n=3) across all timepoints correlated with E/N ratio in same samples (Spearman). (**C**) Lung gene expression by qRT-PCR for *Ccl11*, *Ccl24* and *Ccl26* after low dose (100-300 CFU) aerosol Mtb infection in B6 mice (n=5-12).

**Supplemental Figure S4:**
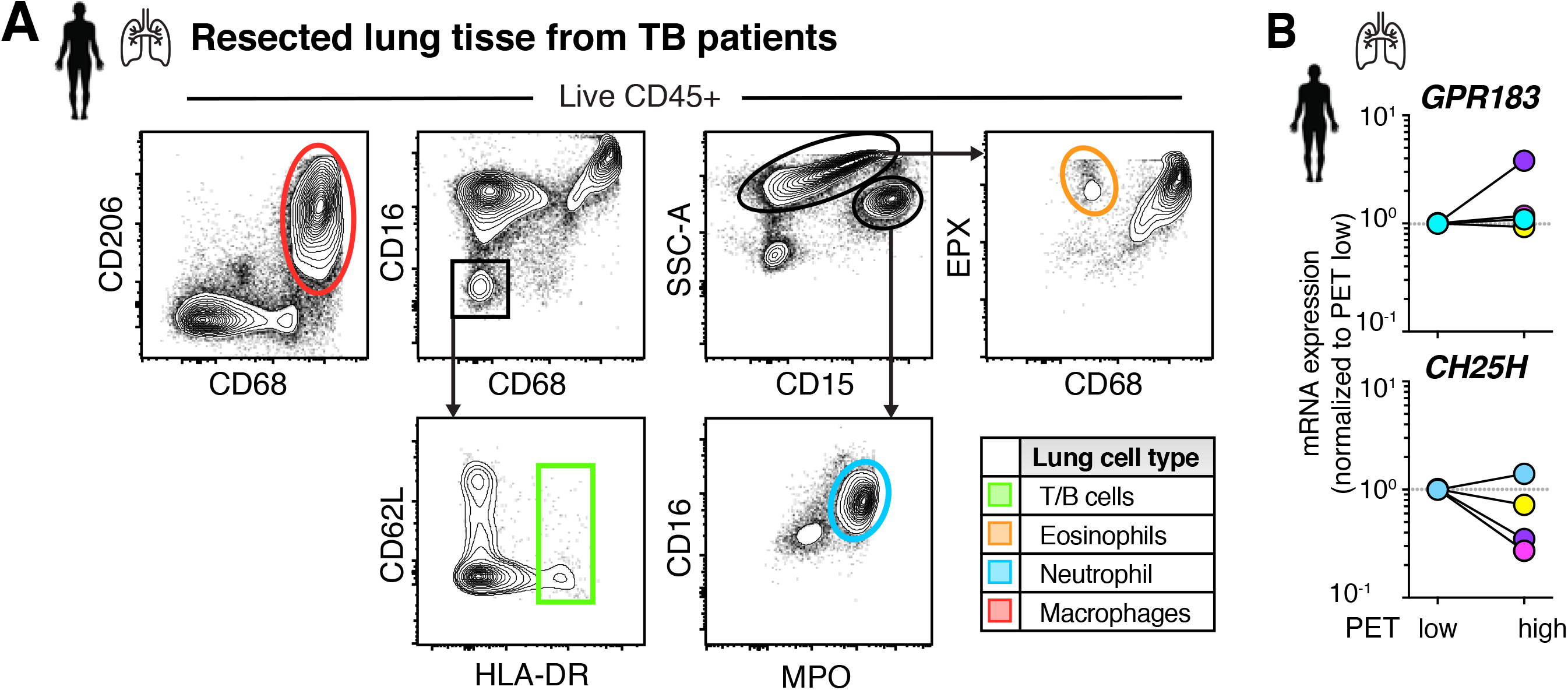
GPR183 expression in human TB lesion. (**A**) Flow cytometric gating strategy of human lung TB lesions for GPR183 expression (n=5) (**B**) qRT-PCR of ^18^FDG PET/CT low or high signal intense human TB lung lesions for *CH25H* AND *GPR183* (n=4, Wilcoxon-matched pairs)

**Supplemental Figure S5:**
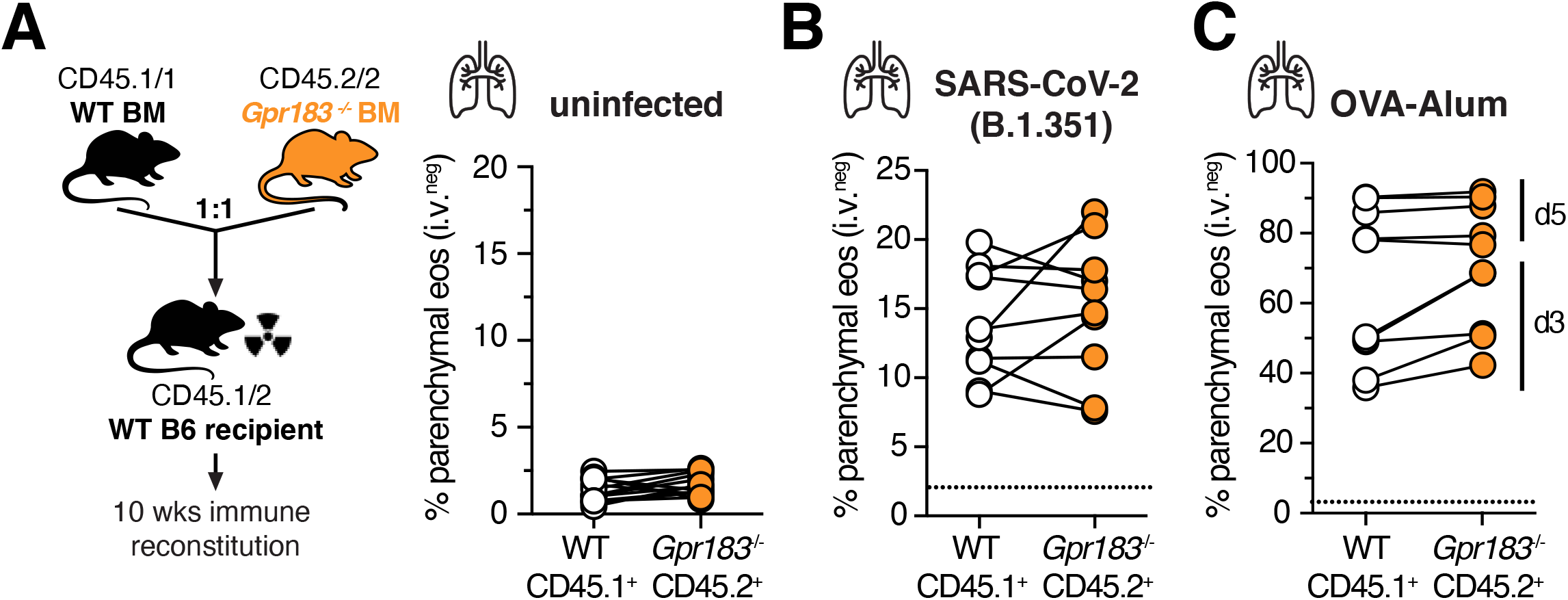
GPR183 expression on eosinophils is dispensable for lung migration in mouse models of viral infection and allergic asthma. (**A**) Frequency of lung parenchymal eosinophils in WT and *Gpr183*^-/-^ competitive mixed BM chimeric mice prior to infection (M, n=13) (**B**) Quantification of CD45 i.v. negative WT or *Gpr183*^- /-^ eosinophils from competitive mixed BM chimeric mice three to five days (F+M, n=9) post intranasal SARS-CoV2 infection (B.1.351, Wilcoxon-matched pairs not significant). (**C**) Quantification of CD45 i.v. negative WT or *Gpr183*^-/-^ eosinophils from competitive mixed BM chimeric mice after OVA-Alum immunization and three (F, n=4) or five days (M, n=5) post intranasal OVA challenge (Wilcoxon-matched pairs not significant, dotted black line shows average CD45 i.v. negative eosinophils or neutrophils in uninfected mixed BM chimeric mice).

**Supplemental Movie: Eosinophil interactions with Mtb and Mtb-infected cells in the airways**

*Ex vivo* lung explant live imaging of d13 Mtb-Cyan Fluorescent protein (CFP) infected EPX-eosinophil reporter mice (pink) with lung structure visualized with anti-laminin (yellow) staining.

## METHODS

### Clinical study population in Shanghai, China

Human TB lung resection samples were obtained at the Shanghai Public Health Clinical Center (SPHCC) between 2015 and 2019 following ethics approval by the SPHCC (# 2015-S046-02 and 2019-S009-02). All patients in the study provided written informed consent, were indicated for lung or pleural tissue resection surgery at SPHCC as an adjunctive therapy for TB (L.W, Y.S.) and were described in Bohrer, Castro et al. (Bohrer et al., 2021).

### Clinical Study population in Bethesda, USA

For whole blood flow cytometry or *in vitro* experiments on eosinophils, healthy blood donor controls from the NIH Clinical Center (Bethesda, MD) were recruited on an institutional review board-approved clinical protocol to obtain blood samples for in vitro research (NCT00001846). All participants gave written informed consent.

### Analysis of publicly available human dataset

We identified nine datasets in the Gene Expression Omnibus (GEO) that profiled whole blood or PBMC samples from patients with LTBI or ATB, of which eight measured *GPR183* expression. After removing samples from patients with other diseases, we analyzed a total of 961 expression profiles across eight datasets: 433 samples from LTBI patients and 528 samples from ATB patients. In addition, we used samples from the Adolescent Cohort Study (ACS - GSE79362), a longitudinal cohort profiling 42 subjects that progressed to active TB and 109 latently infected controls at several time points (prior to diagnosis for progressors, while non-progressors were monitored at regular intervals). We used local regression (LOESS) to model the gene expression profiles over time in each group. We downloaded normalized data from GEO and applied the multi-cohort analysis framework (Haynes et al., 2017) to calculate effect sizes reflecting gene expression differences across clinical groups (latent TB and active TB). We used Hedge’s g as a measure of effect size.

### Experimental animals - Rhesus Macaques

Ten healthy >2-year-old rhesus macaques (male and female, tuberculin skin test negative) were received from the NIAID breeding colony on Morgan Island. Animals were housed in biocontainment racks and maintained in accordance with the Animal Welfare Act, the Guide for the Care and Use of Laboratory Animals and all applicable regulations, standards, and policies in a fully Association for Assessment and Accreditation of Laboratory Animal Care International accredited animal biosafety level 3 vivarium. All procedures were performed utilizing appropriate anesthetics as listed in the NIAID DIR Animal Care and Use Committee approved animal study proposal LPD25E. Euthanasia methods were consistent with the American Veterinary Medical Association guidelines on euthanasia and endpoint criteria listed in the NIAID DIR Animal Care and Use Committee approved animal study proposal.

### Mtb infections of rhesus macaques

Rhesus macaques from three independent infections provided tissues for this study as well as other work reported previously (Bohrer et al., 2021; Namasivayam et al., 2019). Briefly, macaques were infected via bronchoscope instillation of 8-50 colony forming units (CFU) of either Mtb strain H37Rv or mCherry-expressing Erdman into the right lower lung lobe. The infection in these animals was allowed to progress for 8 to 13 weeks with disease progression monitored by PET/CT and physical exam. BAL and blood were collected at indicated timepoints prior to euthanasia and lung tissue collection.

### Experimental animals - Mice

C57BL/6 (B6) mice were purchased from Taconic Farms (Hudson, NY). B6.SJL (CD45.1/1), B6.SJL/ C57BL/6 (CD45.1/2) mice were obtained through a supply contract between the National Institute of Allergy and Infectious Diseases (NIAID) and Taconic Farms. *Epx*-Cre mice (Doyle et al., 2013) were initially obtained from Elizabeth Jacobsen (Mayo Clinic, Phoenix, AZ) and subsequently crossed to ROSA26-LSL-tdTomato mice to generate eosinophil-specific fluorescent EPX-reporter animals and generously gifted by David Sacks (NIAID) (Lee et al., 2020b). *Gpr183^-/-^* mice (Baptista et al., 2019; Liu et al., 2011) were gifted by Michael Fessler (NIEHS) and Vanja Lazarevic (NCI). *Ccr3^-/-^* (JAX 5440, C.129S4-Ccr3tm1Cge/J) on the BALB/c background, BALB/c (JAX 651), BALB/c CD45.1 (JAX 6584, CByJ.SJL(B6)-Ptprca/J) and *Ch25h*^-/-^ (JAX 16263, B6.129S-Ch25htm1Rus/J) mice were purchased from Jackson Laboratories (Bar Harbor, ME). Both male and female mice, 8-24 wk. old at onset of experiments were used and experimental groups in individual experiments were age and sex matched. All animals were bred and maintained in an AAALAC-accredited ABSL2 or ABSL3 facility at the NIH and experiments performed in compliance with an animal study proposal approved by the NIAID Animal Care and Use Committee (protocol LCIM17E).

### Mtb infections of mice

Aerosol infections of mice with H37Rv strains of Mtb were carried out in a whole-body inhalation exposure system (Glas-Col, Terre Haute, IN). Delivery doses (100-300 CFU) were confirmed by measuring lung bacterial loads at 2-24hrs post exposure in control mice. H37Rv-mCherry was kindly provided by Kevin Urdahl (Cohen et al., 2018), and H37Rv-CFP was gifted by Joel Ernst (UCSF).

### Oxysterol measurement by liquid chromatography tandem mass spectrometry

All solvents for oxysterol extraction were LCMS-grade purchased from Thermo Fisher Scientific. Unesterified oxysterols were measured as previously described with modifications (McDonald et al., 2012). BAL fluid was thawed and inactivated via addition of 2 sample volumes of 10 µg/ml butylated hydroxytoluene in LCMS-grade methanol. To 400µl of sample, 10µl of 100µM 27-hydroxycholesterol-d6 (Avanti Polar Lipids) was added as an internal standard and 300µl of standard-containing sample was transferred dropwise to 3ml of a 1:1 dichloromethane:methanol mixture in a nitrogen evacuated glass tube. Samples were incubated for 10min ambient. After incubation, 3ml of dPBS was added. Samples were centrifuged for 10 minutes at 3,800 RPM in a JA-12 rotor (Beckman Coulter Inc.) and the bottom layer collected. The aqueous fraction was reextracted with 3ml of dichloromethane and the organic layers from the first and second extractions were combined. Samples were dried under nitrogen. Oxysterols were enriched by solid phase extraction using 200 mg, 3 mL aminopropyl columns (Waters™). Columns conditioned with 6 mL of hexanes. Dried samples were dissolved in 1ml of hexanes and loaded onto the column. The sample tube was rinsed with an additional 1ml of hexanes which was subsequently loaded to the column. Columns were rinsed with one volume of hexanes. Oxysterols were eluted into test tubes using two column volumes of 23:1 LCMS-grade chloroform:methanol. Samples were dried under nitrogen and resuspended in in 400µl of 90% LCMS-grade methanol and incubated at 37°C for 30 minutes and then prepared for LCMS injection. A 40 µL injection of each sample was separated across a Kinetex® Polar C18 column (100 Å, 2.6 µm, 100 x 3 mm) on a Nexera LC-20ADXR HPLC (Shimadzu). Analytes were eluted at 0.4 ml/min using a binary gradient from (A) 5 mM ammonium acetate in 70% acetonitrile 30% water (v/v) to (B) 5 mM ammonium acetate in 50% isopropanol 50% acetonitrile (v/v) over 7 minutes followed by a 3 min hold at 100 % B. Samples were detected with a 6500+ QTrap mass spectrometer (AB Sciex) using established multiple reaction monitoring pairs and retention times were compared to repeated injections of OxysterolSPLASH™ standard mix (Avanti Polar Lipids) in an extraction control sample of rhesus BAL. All signals were integrated using SciexOS™ 2.0.0 and normalized back to internal standard levels. All statistical analysis was performed in GraphPad™ Prism 8.4.3.

### Confocal microscopy of murine live lung explant sections

Confocal microscopy of murine live lung sections was used to visualize organ architecture, H37Rv-CFP expressing Mtb bacilli and EPX-reported eosinophil cell migration *ex vivo*, at as close to physiological conditions as possible. Briefly, after euthanasia, d13 H37Rv-CFP Mtb-infected EPX-CRE-ROSA26-LSL-tdTomato mouse lungs were inflated with 1.5% low-melt agarose in RPMI at 37°C. Inflated tissues were kept on ice, in 1% FBS in PBS, and sliced into 300 µm sections using a Leica VT1000 S Vibrating Blade Microtome (Leica Microsystems) at speed 5, in ice-cold PBS. Lung sections were then stained with anti-laminin antibody for 2h on ice followed by washing and culturing in complete lymphocyte medium (Phenol Red-free RPMI supplemented with 10% FBS, 25mM HEPES, 50μM β-ME, 1% Pen/Strep/L-Glu and 1% Sodium Pyruvate) in humidified incubator at 37°C where they rested for 12 h prior to imaging. Lung sections were held down with tissue anchors (Warner Instruments) in 14 mm microwell dishes (MatTek) and imaged using Leica SP5 inverted 5 channel confocal microscope equipped with an Environmental Chamber (NIH Division of Scientific Equipment and Instrumentation Services) to maintain 37°C and 5% CO_2_ under BSL3 containment conditions. Microscope configuration was set up for four-dimensional analysis (x,y,z,t) of cell segregation and migration within tissue sections. A Diode laser for 405 nm excitation; an Argon laser for 488 nm and 514 nm excitation, a DPSS laser for 561nm excitation; and HeNe lasers for 594 nm and 633 nm excitation wavelengths were tuned to minimal power (between 1 and 5%). Z stacks of images were collected (10 – 50 µm). Mosaic images of lung sections were generated by acquiring multiple Z stacks using a motorized stage to cover the whole area of the section and assembled into a tiled image using LAS X software (Leica Microsystems). For time-lapse analysis of cell migration, tiled Z-stacks were collected over time (1 to 4h). Post-acquisition mages were processed using Imaris (Bitplane) software.

### Generation of competitive mixed BM chimeric mice

For competitive mixed BM chimeric mice, C57BL/6 B6.SJL (CD45.1/1 or CD45.1/2) wild-type (WT) or BALB/c CD45.1 WT mice were lethally irradiated (950 rad) and reconstituted with a total of 10^7^ donor BM cells from either C57BL/6 or BALB/c CD45.1/1 WT mice mixed in equal parts with BM cells from B6 *Gpr183^-/-^* mice (CD45.2/2) or BALB/c *Ccr3^-/^-* mice (CD45.2/2). Mice were placed on trimethoprim-sulfamethoxazole water for four weeks immediately after irradiation, followed by an additional 4-6 weeks on regular water prior to infection, for a total time allotted for immune reconstitution of 10-12 weeks.

### OVA-Alum lung allergy model in mice

Reconstituted mixed *Gpr183*^-/-^ and WT BM chimeric mice were injected intraperitoneally twice, 14 days apart, with 100µg Ovalbumin (OVA, Sigma-Aldrich, Cat-A5503, Lot-SLBQ9036V) in 200µl containing 12.5% Imject^TM^ alum adjuvant (Thermo Scientific, Cat-77161, Lot-TC260123C). Ten days after the last injection mice were anaesthetized with isoflurane and challenged intranasally with 30µg OVA in 30µl injection grade sterile saline. Lung analyses of eosinophil migration were performed three and five days post intranasal challenge.

### SARS-CoV-2 infection in mice

Reconstituted mixed *Gpr183*^-/-^ and WT BM chimeric mice were anesthetized with isoflurane and infected intranasally with 3.5×10^4^ TCID_50_ SARS-CoV-2 hCoV-19/South Africa/KRISP-K005325/2020 beta variant of concern (Pango lineage B.1.351, GISAID reference: EPI_ISL_860618, BEI resources) in 30µl. Viral inoculum was quantified by TCID_50_ assay in Vero E6 cells (CRL-1586; American Type Culture Collection). Lung analyses of eosinophil migration were performed three and five days post intranasal challenge. Viral stocks were generated by infection of VeroE6 cells stably expressing TMPRSS2 (Liu et al., 2021) at a multiplicity of infection of 0.01 for 48hrs. Cell culture media was harvested and centrifuged at 3500xg, pooled, aliquoted and stored at −80°C until use. Virus stock was sequenced using the Illumina platform and contained the single nucleotide variants consistent with the reference sequence.

### RNA extraction, cDNA synthesis, and quantitative real-time polymerase chain reaction (qRT-PCR)

For RNA extraction from murine lung tissue dedicated lung lobes were placed in RNAlater (Invitrogen, Carlsbad, CA) and stored at −80°C for later analysis. RNAlater-stabilized lung lobes were thawed at RT for 20min, then homogenized in RLT plus buffer with β (Qiagen, Hilden, Germany). Total RNA was then isolated from the RLT-homogenized cells using the RNeasy Plus Mini Kit (Qiagen, Hilden, Germany). Individual lung RNA samples and FACS purified AM samples were reverse-transcribed using Superscript IV First-Stand Synthesis System (ThermoFisher Scientific, Waltham, MA) according to the manufacturer’s instructions using random hexamers. Quantitative RT-PCR was performed in a CFX384 Optical Reaction Module (Bio-Rad Laboratories, Hercules, CA). Relative quantities of mRNA for several genes were determined using iTaq Universal SYBR Green Supermix (Bio-Rad Laboratories, Hercules, CA) normalized to *ActB* mRNA levels and then expressed as a relative increase or decrease in arbitrary units (AU) or as fold changes over indicated biological controls. Primers were designed using Roche Universal Probe Library (Roche, Basel, Switzerland). The sequences of the specific primers are as follows: *Ccl11:* agagctccacagcgcttct, gcaggaagttgggatgga; *Ccl24:* gcagcatctgtcccaagg, gcagcttggggtcagtaca; *Ccl26:* gttcttcgatttgggtctcct, ggacatagcgatgctgctc; *Ch25h:* acccactacccatattta; gcccagcattttgtccca; *ActB:* ctaaggccaaccgtgaaaag; accagaggcatacagggaca.

For RNA extraction from human lung resection material 50∼100 mg of RNAlater-stabilized lung tissue pieces were added to 1ml of Trizol (Invitrogen) and homogenized. Chloroform (0.2ml) was added and incubated for 3min. After centrifugation for 15min at 4°C, the aqueous phase was transferred to a new 1.5ml tube. Isopropanol (0.5ml) was added and incubated for 10min. After centrifugation for 10min at 4°C, the supernatant was discarded, and the pellet re-suspended in 1ml of 75% ethanol. After vortexing and centrifugation, the RNA was pellet air-dried for 10min and re-suspended in 20μl RNase-free water. Total RNA was then subjected to genomic DNA (gDNA) elimination and reverse-transcription according to PrimeScript™ RT reagent kit protocols (Takara). Briefly, total RNA was incubated with genomic DNA eraser for 2min at 42°C (total volume of 10μl containing 2μl of 5x gDNA eraser buffer, 1μl of gDNA eraser, 1μg of total RNA and RNase-free water). Next RNA was incubated with reverse-transcription enzyme mix (total volume of 20μl containing 10μl of solution from the above step, 4μl of 5x primescript buffer, 1μl of primescript RT enzyme mix I, 1μl of RT prime mix and RNase-free water) for 15min at 37°C, followed by 5sec at 85°C to synthesize cDNA. Another 80μl of RNase-free water was added before its storage. The cDNA was subjected to RT-PCR by using the Hieff qPCR SYBR Green Master Mix (Shanghai Yeasen Bio Technologies) according to the manufacturer’s instructions. Briefly, the cDNA was incubated with the primers and SYBR Green mix (total volume of 50μl containing 25μl of SYBR green master mix, 1μl of forward primer, 1μl of reverse primer, appropriate volume of cDNA and RNase-free water), and run on a thermal cycling system (CFX96, Bio-Rad) as follows: hot-start at 95°C for 5min, and 40 cycles of 95°C for 10 sec, 60°C for 30 sec, followed by a default stage of melting curve analysis. The quantitative evaluation was performed according to the fusion fluorescence signal. The sequences of the specific primers are as follows: *CCR3*: AGCCTTCCACACTCACCTCTA, ATTAAGCAGGGAAAGAACTAGGC *CCL11*: CCCCAGAAAGCTGTGATCTTCAA, AACCCATGCCCTTTGGACTG, *CCL2*4: GCAGGAGTGATCTTCACCAC, CATCACTGTGGCTTCTCCAGG; *CCL26*: CCCAGCGGGCTGTGATATTC, ACTCTGGGAGGAAACACCCT; *CH25H*: CACCCTGACTTCTCGCCATC, CTTGTGGAAGGTGCGGTACA, *GPR183*: CGAGTCACTGATATACACCTGGA, AAGTTTCCCACGAGCCCAAT

### Cell Isolations for flow cytometry

Rhesus macaque and human peripheral blood samples were collected in EDTA tubes and whole blood was used for flow cytometry. Briefly, antibody-cocktails were directly added to 200μl of whole blood and stained at 37°C for 20min, after which samples were washed twice in 4ml 1% FCS/PBS. Fixable live/dead cell stain (Molecular Probes-Invitrogen) was used according to the manufacturer’s protocol. Pellets were then fixed, permeabilized and red cells lysed with the eBioscience Transcription Factor Staining Buffer Kit (Life Technologies/eBioscience) for at least 1hr followed by intracellular antigen staining for 30min at 4°C. For murine whole blood, 50μl of blood samples (retro-orbital bleed) were collected in EDTA tubes and washed twice immediately with 1ml of 1% FCS/PBS followed by staining with antibody-cocktails for 20min at 4°C. Fixable live/dead cell stain (Molecular Probes-Invitrogen) was used according to the manufacturer’s protocol. Pellets were then fixed, permeabilized and red cells lysed with the eBioscience Transcription Factor Staining Buffer Kit (Life Technologies/eBioscience) for at least 1hr followed by intracellular antigen staining for 30 minutes at 4°C. Rhesus macaque BAL samples were passed through a 100μm cell strainer, pelleted, and counted for analysis. Samples were then stained as described above. The surgically resected human lung tissues were processed and stained as recently described in detail (Bohrer et al., 2021).

Mice were intravenously (i.v.) injected 3min prior to euthanasia with 5-8μg per mouse of APC or BV711 labelled CD45 (30-F11) as previously reported (Anderson et al., 2014). Lungs from infected mice were dissociated via GentleMACS (Miltenyi Biotec, CA) in digestion buffer comprised of Miltenyi lung cell isolation buffer (Miltenyi Biotec, CA), benzonase (7U/ml, Sino Biological), cytochalasin D (10μM, Sigma-Aldrich) and hyaluronidase (0.2mg/ml, Sigma-Aldrich) followed by 30-45 minutes at 37°C. Digested lung was fully dispersed by passage through a 100μm pore size cell strainer and an aliquot removed for bacterial load measurements when needed. Isolated cells were stained as described above. Samples were acquired on a FACSymphony (BD Biosciences) at NIH or on a LSR-Fortessa (BD Biosciences) at SPHCC. FACS data were analyzed using FlowJo10 (Treestar, Ashland, OR).

### FACS sorting of Mtb-infected AM from BAL

Bronchoalveolar lavages (BAL) with 5x 1ml of 0.1mM EDTA PBS were performed 14 days post Mtb infection in WT B6 mice and recovered BAL cells from 7 mice were pooled for each experiment. Bronchoalveolar lavage cells were then washed with cRPMI without phenol red, counted and subsequently surface stained under sterile conditions. AM were sorted based on Mtb fluorescence into infected Mtb (H37Rv)-mCherry^+^ AM or uninfected bystander Mtb (H37Rv)-mCherry^neg^ AM after gating on live CD45.2^+^, CD11c^+^, Siglec-F^+^ cells. Samples were sorted on a FACS Aria III (BD Biosciences, San Jose, CA) under BSL3 conditions and analyzed using FlowJo10 Software for Mac (Treestar, Ashland, OR). Cells were washed with sterile PBS and total RNA was isolated immediately after sorting using the RNeasy Plus Micro Kit from Qiagen (Hilden, Germany) according to the manufacturer’s protocol.

### Human eosinophil isolation and transwell migration assay

Granulocyte layers were isolated from peripheral blood after density gradient centrifugation (Ficoll-Paque PLUS, GE Healthcare, Uppsala, Sweden) and used for eosinophil purifications by negative selection on an AutoMACS Pro separator using the Eosinophil Purification Kit (Cat. No. 130-092-010, Miltenyi Biotec, Bergisch Gladbach, Germany). Eosinophil purity was >98% for all samples as determined by counting a minimum of 300 cells on cytospin slides stained with Shandon™ Diff-Kwik™ stains (Cat. No. 9990700, ThermoFisher Scientific, Waltham, MA).

Transwell migration assays were performed in 24-well Transwell plates with 5.0 μm pore inserts (Costar, Corning, NY). 500 μl of phenol-red free RPMI medium (10%FCS), with or without 100 ng/ml recombinant human CCL11 (eotaxin-1, Cat. No. 300-21, Peprotech, Cranbury, NJ), 100nM 25-OHC (Cayman Biochemicals), 100nM 7α, 25-DHC (Cayman Biochemicals) or the GPR183-specific antagonist NIBR189 (50nM) (Tocris) was added to the lower chamber. 100μl of culture medium containing 3 x 10^5^ cells were then added to the upper chamber. After incubation for 3 hours at 37 °C and 5% CO_2_, cells were collected separately from the upper or lower chambers and acquired on a LSRII flow cytometer (BD). The proportion of migrating eosinophils was defined as the number of cells in the lower chamber divided by the sum of the cells in the upper and lower chambers and normalized to the positive control of eotaxin-mediated migration.

### Multiplex protein analysis in rhesus BAL

For rhesus macaque BAL samples were sterile filtered and CCL11 (Cat. No. HCYTOMAG-60K, MilliporeSigma, Burlington, MA), CCL24 and CCL26 (Cat. No. HCYP2MAG-62K, MilliporeSigma, Burlington, MA) and EPX and MBP were measured in supernatants by suspension array multiplex immunoassays as described previously (Makiya et al., 2014).

### Flow cytometry reagents

Fluorochrome-labeled antibodies against mouse, human or rhesus macaque antigens used for flow cytometric analysis are listed below:

I-A/I-E (clone M5/114.15.2), Ly6G (1A8), CD11c (HL3 and N418), CD45.1 (A20), CD45.2 (104), CD11b (M1/70), CD45 (30-F11), CD68 (FA-11), Ly6C (AL-21 and HK1.4), CX3CR1 (SA011F11), Siglec-F (E50-2440), CD193 (J073E5), CD193 (5E8.4), EPX (AHE-1), CD62L (MEL-14), CD294 (BM16), CD117 (104D2), CD14 (M5E2), CD11c (3.9), CD45 (D058-1283), CD66abce (TET2), CD15 (HI98), CD68 (Y1/82A), HLA-DR (L243), CD16 (3G8), MPO (MPO45S-8E6), CD45 (2D1), CD193 (83101), GPR183 (SA313E4), CD66b (G10F5), CD62L (REG-56), CD206 (15-2.2), CD123 (6H6) and Siglec-8 (7C9).

### Statistical analyses

The statistical significance of differences between data groups was calculated using GraphPad Prism 8 and the Mann-Whitney test, the Wilcoxon matched pairs test (mixed BM chimera experiments, PET -low vs high or otherwise paired samples and Spearman test for correlations were utilized unless otherwise indicated in the figure or figure legend.

